# Spatiotemporal dissociation of brain network connectivity in salience processing: A simultaneous pupillometry-EEG-fMRI study

**DOI:** 10.1101/2022.01.29.478345

**Authors:** Hengda He, Linbi Hong, Paul Sajda

**Affiliations:** Department of Biomedical Engineering, Columbia University, New York, NY 10027; Department of Electrical Engineering, Columbia University, New York, NY 10027; Department of Radiology, Columbia University, New York, NY 10032; Data Science Institute, Columbia University, New York, NY 10027

## Abstract

The processing of salient stimuli involves a wide range of both bottom-up and top-down processes. Previous neuroimaging studies have identified multiple brain areas and networks for salience processing, including the salience network (SN), dorsal attention network (DAN), and the locus coeruleus-norepinephrine (LC-NE) neuromodulatory system. However, interactions among these networks and the cortico-subcortical systems in salience processing remain unclear. Here, we simultaneously recorded pupillometry, electroencephalogram (EEG), and functional magnetic resonance imaging (fMRI) during an auditory oddball paradigm. Using EEG-informed fMRI analysis, we temporally dissociated the target stimulus evoked activation, allowing us to identify the cascades of cortical areas associated with salience processing. Furthermore, functional connectivity analysis uncovered spatiotemporal functional network organizations of these salience processing neural correlates. Using pupillometry as a psychophysiological marker of LC-NE activity, we also assessed brain-pupil relationships. With state-space modeling of target modulated effective connectivity, we found that the target evoked pupillary response is associated with the network causal couplings from late to early subsystems, as well as the network switching initiated by the SN. These findings indicate that the SN might cooperate with pupil-indexed brainstem neuromodulatory systems, such as the LC-NE system, in the reorganization and dynamic switching of cortical networks, and shed light on the implications of their integrative framework in various cognitive processes and neurological diseases.

## Introduction

To navigate complex and dynamic environments our brains cannot allocate attention to everything, but instead must continuously mark and process salient objects [97, 98]. For example, when we are walking on a busy street, we will likely direct attention to the traffic lights, a siren, and our planned route. In psychology and neuroscience, the term ‘salience’ refers to a noticeable or important object that stands out from the surroundings or background. Salience is usually accompanied by unexpectedness, novelty and infrequency [42]. Typically, salience processing involves two general mechanisms [63]: 1) bottom-up processing that includes filtering and amplifying the sensory information; 2) top-down processing in support of anticipation, cognitive control and goal-directed behaviors. To investigate salience processing, one of the widely used experimental paradigms is the oddball task, where subjects are instructed to detect distinct infrequent target stimuli in a stream of standard stimuli. In previous functional MRI (fMRI) studies, a variety of brain areas have been identified as correlates of salience processing, including regions in the ventral and dorsal attention networks (VAN and DAN), salience network (SN), sensory cortex, primary somatosensory cortex (S1), subcortex [42, 48, 49]. However, it is challenging to dissociate and interpret the distinct cognitive processes underlying these spatially distributed regions. Even though functional connectivity analyses have been used to dissociate brain networks [83, 88], the lack of time scales and the directionality in the coupling of these brain regions and networks still hinder the inference of their roles in salience processing.

Besides cortical networks, the locus coeruleus (LC), as the primary source of norepinephrine (NE), has also been associated with salience processing. The phasic LC activity has been shown to produce the P300 event-related potential (ERP), which typically appears robustly following target stimuli (responds weaker following standard stimuli) in oddball paradigms [3, 105]. Besides the P300 ERP, pupil diameter has also been used as a psychophysiological marker of the LC activity [68]. For example, in a single-unit recording study, both the spiking activity in the LC and pupil diameter are evoked following unexpected auditory stimuli [47]. Trial-by-trial associations were also observed between the pupillary response magnitude and LC responses. The association between the activity in LC and pupil diameter fluctuations has also been shown in a fMRI study with oddball paradigm [67]. Together, these findings indicate the reliability of LC-pupil relationships during neural processes of salience stimuli in the oddball paradigm. Pupil diameter fluctuations reflect salience, attention, surprise, efforts and arousal [46]. In the oddball paradigm, target-driven pupil dilation reflects both bottom-up adaptation and neural selectivity for stimulus features, and top-down cognitive processes of decision-making and task demands [46].

Both the cortical network dynamics and the LC-NE system have been well characterized, such as the triple-network switching model of the SN [62], and the adaptive gain theory of the LC [3, 35]. Even though the SN and the LC-NE system have been closely related to each other [3, 22, 55, 57] and both centrally positioned in attentional processing, it is still unclear what their integrated roles are in salience processing. Hence, it would be valuable to investigate the cortico-subcortical interactions between the cortical network dynamics and pupil-indexed neuromodulatory systems, such as the LC-NE system. Critically, a better understanding requires the assessment of the causal couplings between cortical networks and brain-pupil relationships in the context of salience processing.

Emerging evidence from the recent literature indicates that neuromodulatory systems, such as the LC-NE system, are important factors in shaping functional network connectivity, reorganization, and dynamics [39, 84, 102, 103, 110]. Thus, in this study, we explored this possibility using simultaneous recordings of pupillometry, electroencephalography (EEG), and fMRI in an oddball paradigm. We first used a single-trial variability (STV) EEG-informed fMRI analysis, which allowed us to map the neural cascade underlying salience processing. Second, with the functional connectivity analyses of the fMRI data, we were able to identify dissociable spatiotemporal functional network organizations of these neural correlates. Then, by leveraging the temporal dynamics of EEG, we further characterized the causal interactions between these regions with a state-space model [96]. Finally, we assessed brain-pupil relationships, which indicate the cortico-subcortical interactions between the cortical network dynamics and pupil-indexed LC-NE system. Specifically, we hypothesized that the pupil-indexed LC activity is associated with the SN dynamic switching model. Our results suggest an integrated role of the SN and the LC-NE system in salience processing.

## Results

### Pupillometry Analysis

We used a MRI-compatible eye-tracking camera to track pupil diameter fluctuations in parallel to simultaneous recordings of EEG and fMRI which measured brain activity. Pupil diameter data were then preprocessed and epoched (see SI Methods for details). The stimuli-locked grand average pupil diameter fluctuations are shown in Fig. S1. We observed a slow pupil dilation evoked by the target oddball stimuli peaking around 1.4 s after the stimulus. To quantify the pupil dilation elicited by the salience stimuli, we extracted the maximum percentage pupil diameter change within each trial as task-evoked pupillary response (TEPR).

### Single-trial EEG-informed fMRI Analysis

To identify the neural correlates involved in the salience processing cascade, we performed a single-trial EEG-informed fMRI analysis. Briefly, this method extracts the EEG components discriminating target versus standard trials at different temporal windows spanning the trial, and these are used to map the temporal evolved brain activities that are correlates of salience processing using the fMRI data (see SI Methods for details). From the resulting group-level whole-brain blood-oxygen-level-dependent (BOLD) activation maps, we identified significant clusters (p < 0.05, cluster corrected) at specific windows as shown in Fig. 1A. These results revealed brain regions associated with salience stimuli processing: left superior parietal lobule (lSPL) (225 ms; positive cluster), left S1 (lS1) (250, 275, 350 and 375 ms; positive cluster), left orbitofrontal cortex (lOFC) (375 ms; negative cluster), left inferior parietal lobule (lIPL) (375 ms; negative cluster), left auditory cortex (lAuditory) and frontal operculum (600 ms; negative cluster), right primary motor cortex (rM1) (225 ms; positive cluster), right secondary visual cortex (rV2) (275 ms; positive cluster), right SPL (rSPL) (275 and 300 ms; positive cluster), right OFC (rOFC) and inferior frontal cortex (rIFC) (375 ms; negative cluster), and supplementary motor area (SMA) and medial prefrontal cortex (mPFC) (400, 425, and 600 ms; negative cluster). These results indicate a coordinated task-related neural cascade, representing the spatiotemporal dynamics of the neural correlates in salience processing.

**Figure 1.**
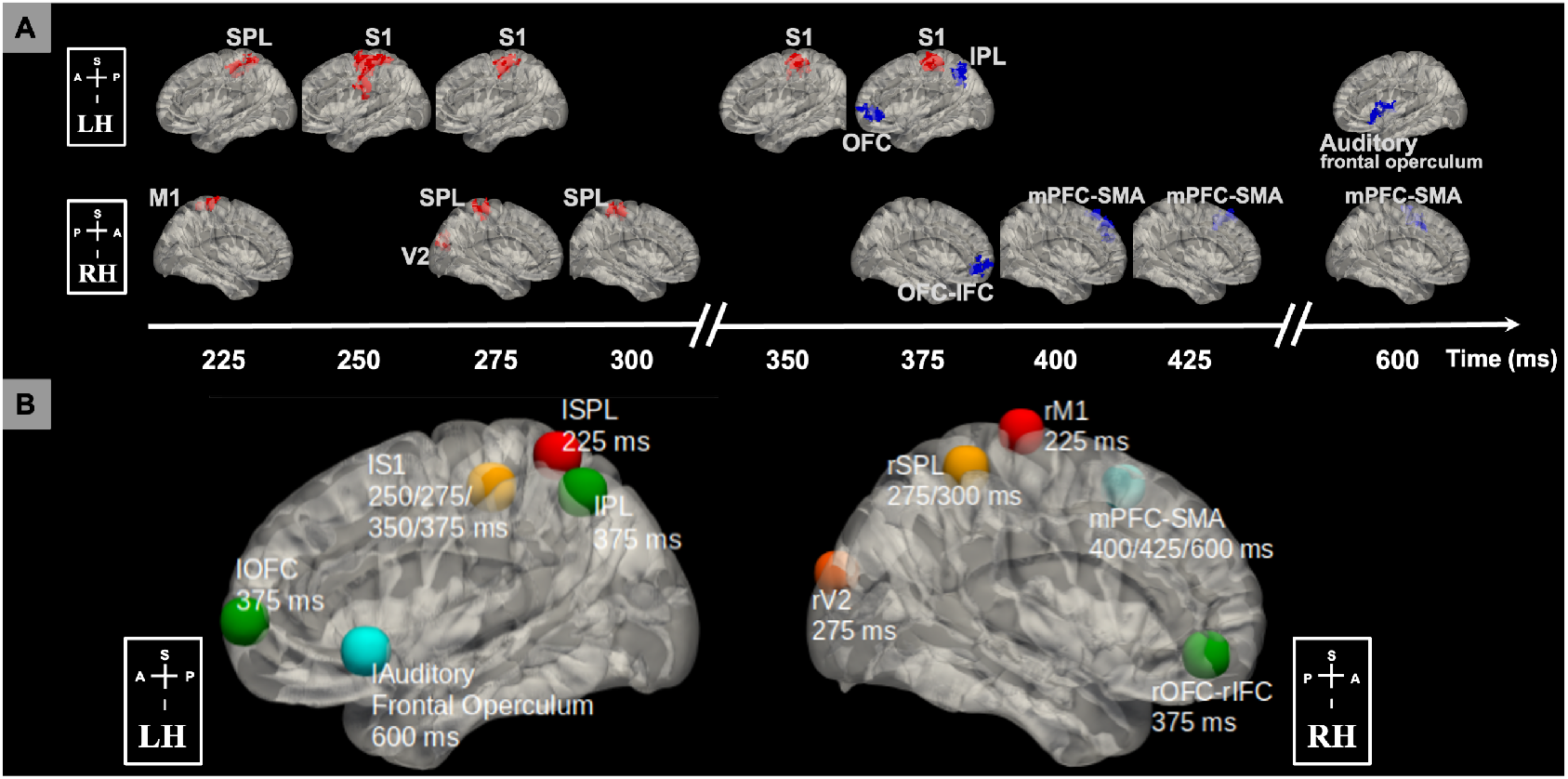
Neural correlates of salience processing defined with the EEG single-trial variability (STV) informed fMRI analysis. (A) Timing diagram showing significant group-level activation clusters (p < 0.05 cluster-wise multiple comparison correction). STV in EEG temporal components discriminating the target versus standard trials was used to map the spatiotemporally distributed BOLD fMRI correlates spanning the trial. EEG STV information was incorporated as BOLD predictors in voxel-wise general linear model (GLM) analysis of fMRI, controlling for the variance due to the presence of stimuli and response time (RT). Cluster colors denote positive (red) and negative (blue) effects. Time is relative to stimulus onset. (B) Definition of salience processing nodes. Each node is a sphere centered on the peak voxel of the group-level STV EEG-informed fMRI analysis results. Centroid of peak locations was used for regions involved in more than one temporal windows. Node colors denote timing of involvement in the trial from early to late (temporal order: red, orange, yellow, green, and blue). All clusters and nodes were overlaid on a 3D Montreal Neurological Institute (MNI) 152 brain pial surface for visualization. BOLD, blood-oxygen-level-dependent; RH, right hemisphere; LH, left hemisphere; A, anterior; P, posterior; S, superior; I inferior; SPL, superior parietal lobule; M1, primary motor cortex; S1, primary somatosensory cortex; V2, secondary visual cortex; OFC, orbitofrontal cortex; IPL, inferior parietal lobule; IFC, inferior frontal cortex; mPFC, medial prefrontal cortex; SMA, supplementary motor area.

### Network Organization of Brain Regions Associated with Salience Processing

Following this observed neural cascade associated with salience processing, a natural question we asked was about the organization of these spatiotemporally distributed regions. Specifically, we aimed to assess the network organization and connectivity between these brain regions. Thus, we defined 10 nodes: lSPL [x = −34, y = −52, z = 64; Montreal Neurological Institute (MNI) coordinates], lS1 (x = −46, y = −28, z = 52), lOFC (x = −40, y = 60, z = 4), lIPL (x = −48, y = −60, z = 50), lAuditory and frontal operculum (x = −54, y = 16, z = −6), rM1 (x = 18, y = −22, z = 76), rV2 (x = 8, y = −94, z = 22), rSPL (x = 38, y = −42, z = 60), rOFC-rIFC (x = 42, y = 46, z = −8), mPFC-SMA (x = 4, y = 18, z = 56) as shown in Fig. 1B.

Given the emerging evidence that indicates the relevance between task activation and the intrinsic network organization of the brain [19, 45], we hypothesized that the previously identified nodes might represent organized underlying brain networks involved in salience processing. To test this hypothesis, a network localization approach was performed to map the brain regions functionally connected with each node (see SI Methods for details; results in Fig. 2). As expected, node rM1 was localized within the sensory motor network, and rV2 was part of the visual network. Intriguingly, lSPL, rSPL and lS1, which are spatiotemporally heterogeneous regions in the salience processing cascade, were mapped to a single brain network, i.e. DAN. Similarly, lOFC, rOFC-rIFC and lIPL, which are all correlated with the EEG discriminating components at 375 ms post-stimulus, fell within the executive control network (ECN). We found the nodes correlated with the late discriminating components, i.e. mPFC-SMA, lAuditory and left frontal operculum, were part of the SN. Overall, this observation suggests a spatially organized intrinsic brain network of these distributed nodes, indicating that the temporal evolution of different task activation regions (Fig. 1A) spanning the trial might be supported by a specific set of brain networks (Fig. 2).

**Figure 2.**
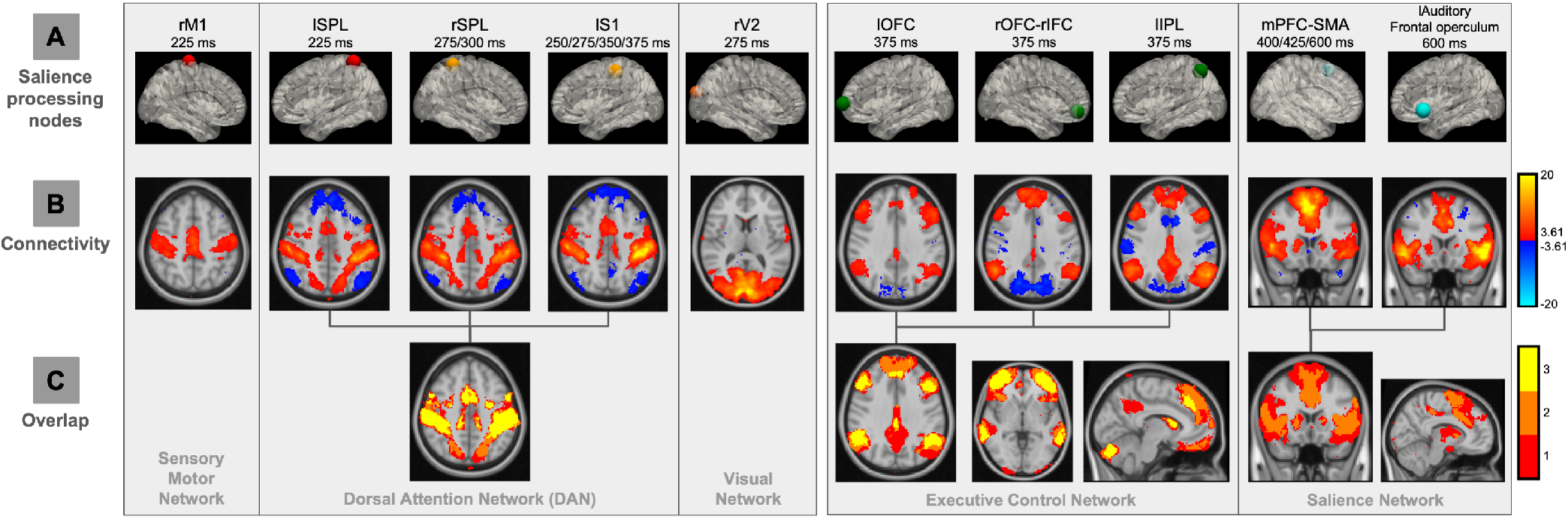
Network localization approach to map functional networks underlying salience processing nodes. (A) BOLD signals from the nodes (intersected with the gray matter mask) were extracted, controlling the nuisance signals (motion-related, ventricle and white matter signals). (B) Group-level functional connectivity (FC) results of each node (t-value, mixed-effect, p < 0.001 uncorrected). Seed-based FC analysis (with the task-related variability regressed out) was used to map network of regions connected to each node location. Colors denote positive (red) and negative (blue) correlations. (C) Spatial overlaps in the FC maps of each node identified spatial network organizations of salience processing nodes. Colors represent the number of FC maps overlapped. lSPL and rSPL, left and right superior parietal lobule; rM1, right primary motor cortex; lS1, left primary somatosensory cortex; rV2, right secondary visual area; lOFC and rOFC, left and right orbitofrontal cortex; lIPL, left inferior parietal lobule; rIFC, right inferior frontal cortex; lAuditory, left auditory cortex.

Following these results that distinct nodes might fall within a common network, our next objective was to directly examine the connectivity between the nodes. As shown in Fig. 3 (p < 0.05, uncorrected), in line with the previously observed spatial organization of the nodes revealed by network connectivity (Fig. 2), we found strong connections across the nodes within each network. For example, lSPL, rSPL and lS1 showed a stronger within network (i.e. nodes of DAN) connectivity compared to their connections with other nodes. Furthermore, the functional connectivity results clearly identified three distinct groups of the nodes, organized by the EEG discriminating component time windows, indicating a temporal organization of the nodes. Thus, to assess the brain networks involved in the task-related neural cascade, we defined three intrinsically connected salience processing networks, i.e. early-time (nodes: lSPL, rM1, rV2, rSPL and lS1), middle-time (nodes: lIPL, lOFC and rOFC-rIFC), and late-time (nodes: mPFC-SMA, lAuditory and left frontal operculum) networks.

**Figure 3.**
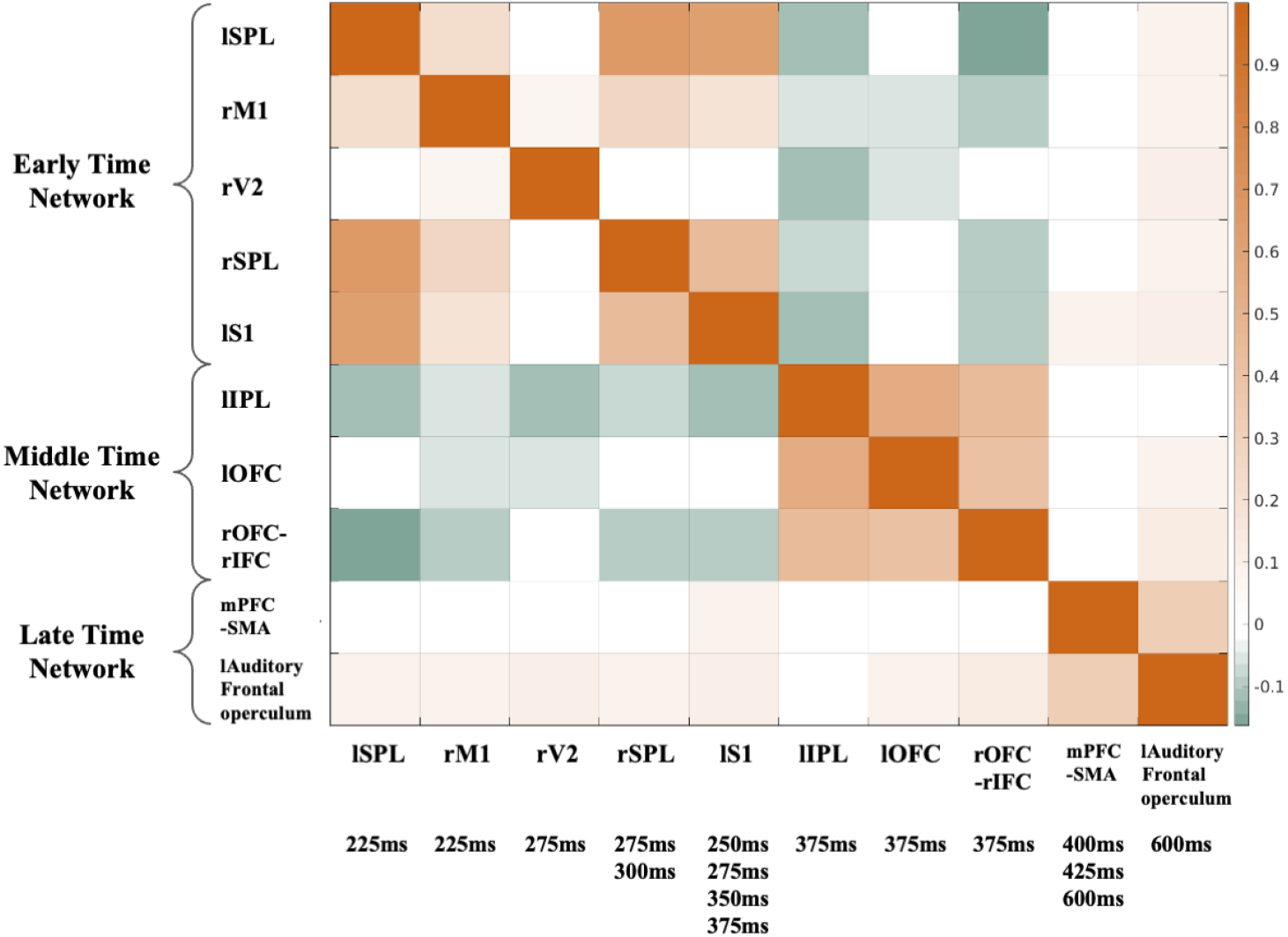
Functional connectivity (FC) across salience processing nodes (group averaged z-score, mixed-effect, p < 0.05 uncorrected). BOLD signals from the nodes (intersected with the gray matter mask) were extracted, controlling the nuisance signals (motion-related, ventricle and white matter signals). Pearson’s correlation was calculated between BOLD signals from the nodes (with the task-related variability regressed out). FC results identified three distinct groups of the nodes, organized by the EEG discriminating component time windows, indicating a temporal network organization of the nodes: 1) early-time network includes lSPL and rSPL, rM1, rV2, and lS1; 2) middle-time network includes lOFC and rOFC, lIPL, and rIFC; 3) late-time network includes mPFC, SMA, lAuditory, and left frontal operculum.

### Modulated Effective Connectivity by Salience Processing

To investigate the directed causal interaction modulated by the salience stimuli, EEG data were fit with a state-space model to infer the effective connectivity (EC) between the nodes (see SI Methods for details). Results for the significant group-level mean EC (Bayesian parameter averaging; *α* < 0.05; Bonferroni corrected) are shown in Fig. S2, with positive and negative connections shown in Fig. 4. To quantify the connection strength of each node, we computed the total connection strength (Fig. S3), which is the sum of all the unsigned connection parameters (efferent, afferent and self-connection) associated with the node. These results suggest that the lSPL and mPFC-SMA are the hubs in the processing of salience stimuli.

**Figure 4.**
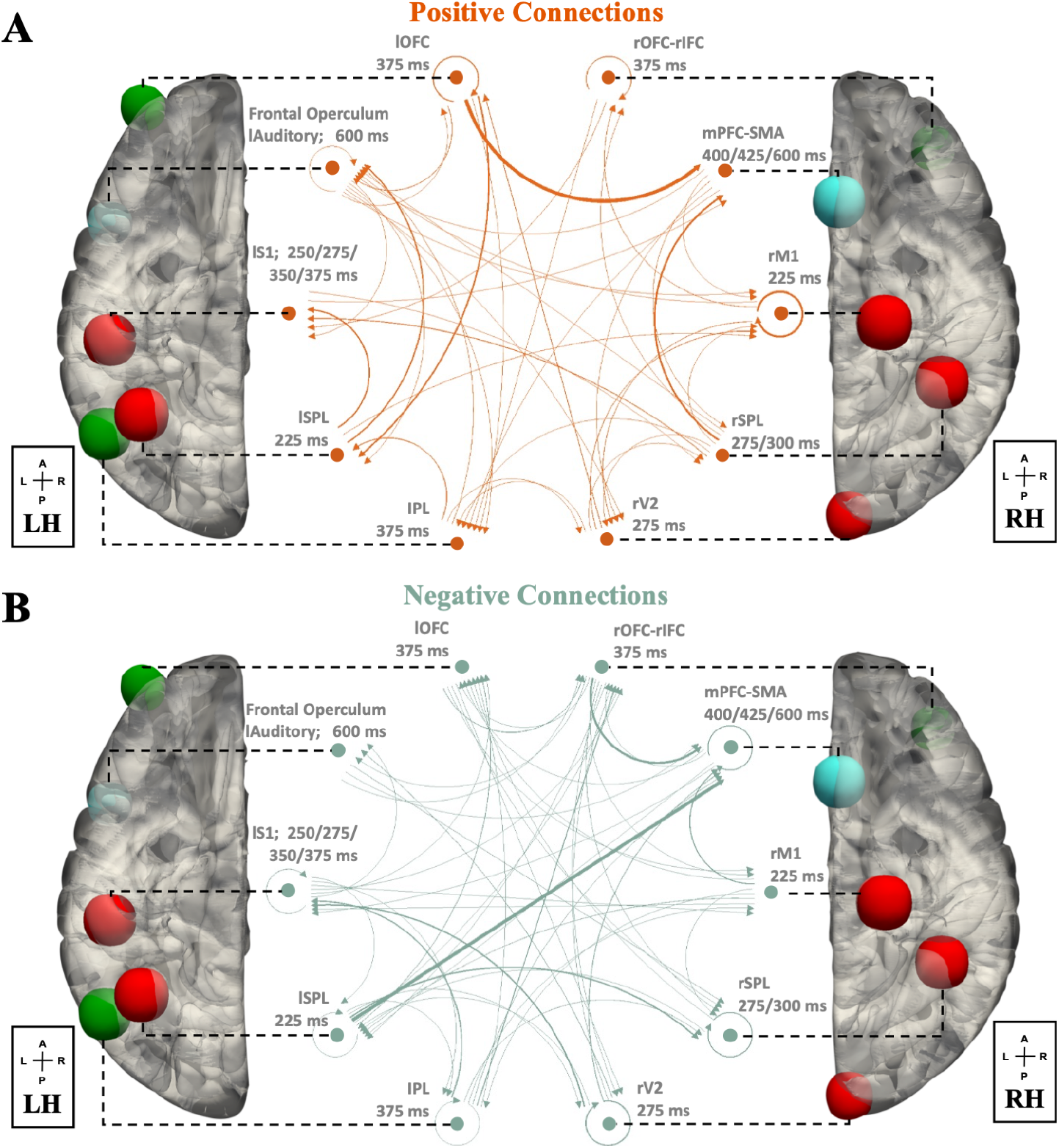
Effective connectivity (EC) across salience processing nodes (Bayesian parameter averaging, *α* < 0.05, Bonferroni corrected). (A) positive EC, (B) negative EC. The arrow and thickness of the connecting lines correspond to the directionality and strength of EC, respectively. Dominant influence is observed in the connections of lSPL, lOFC and mPFC-SMA. Node colors denote timing of involvement (early-time: red; middle-time: green; late-time: blue).

### Effective Connectivity of Networks Predicts Pupillary Response

Having demonstrated the causal interaction between the nodes modulated by the salience stimuli, we next investigated the network-level EC, based on previously defined salience processing networks (Fig. 3). To characterize the network-level connectivity in the positive and negative connections (Fig. 4), we computed positive and negative network strength as the sum of all positive and negative connection parameters from one network node set to the other network node set (or to itself as self-connection network strength), respectively. The network strength of group-level mean EC is shown in Fig. S2.

We next assessed the relevance of network EC strength to TEPR, by computing the Pearson correlation between positive (or negative) network strength and TEPR at the between-subject level. We found a significant correlation between the late-to-early positive network strength and TEPR (r = 0.6352, p = 0.0035; Fig. 5A; after controlling response time (RT) and model evidence lower bound (ELBO): r = 0.6347, p = 0.0035), whereas other network interactions did not show a significant correlation (Bonferroni correction; SI Results). This outcome suggests that the early-time network receives salience stimuli modulated excitatory feedback from the late-time network, and this network coupling may be associated with the TEPR.

**Figure 5.**
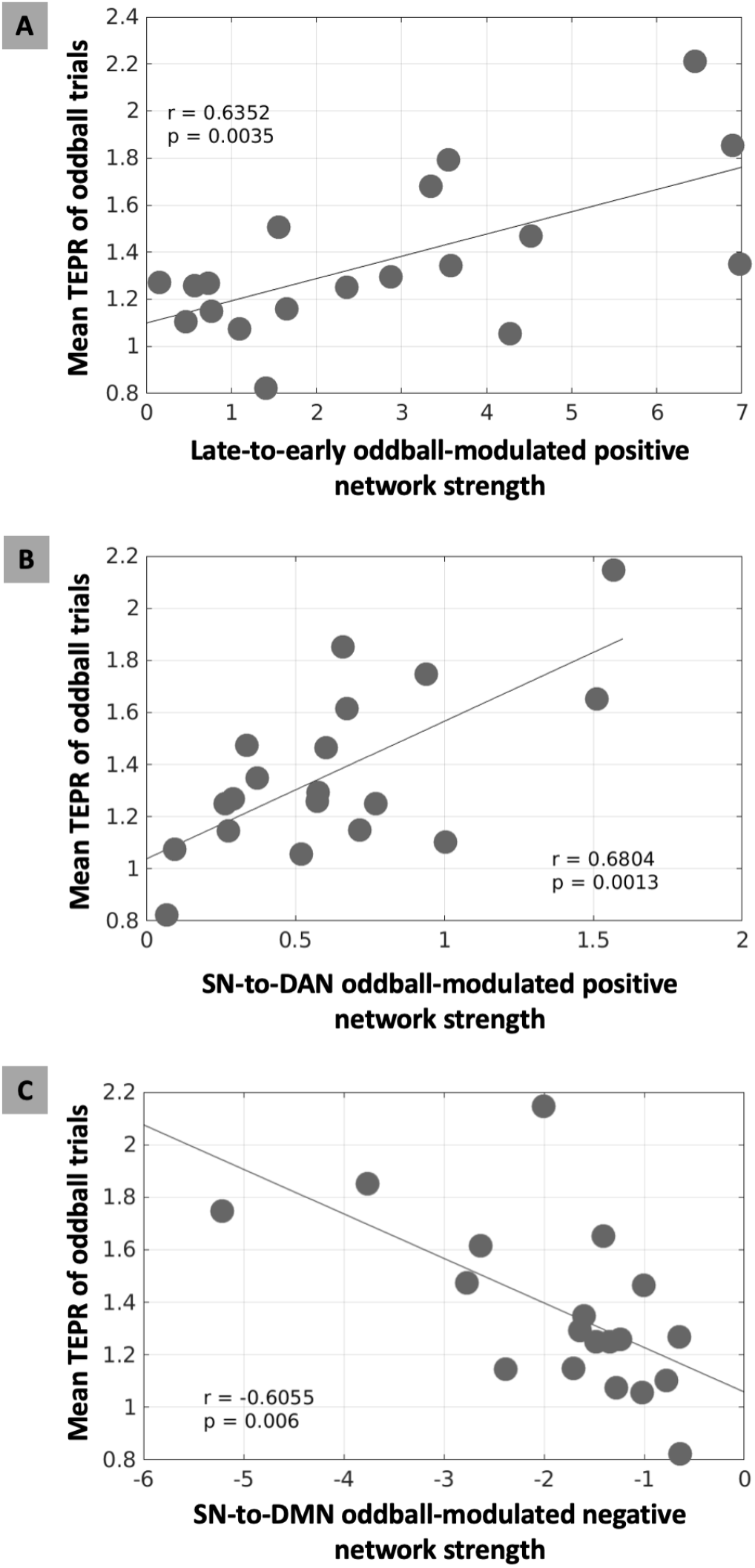
Brain-pupil relationships of the cortical network-level effective connectivity (EC) and task-evoked pupillary response (TEPR) in salience processing. (A) The oddball-modulated positive EC strength from the late-time to early-time network correlated with higher TEPR of oddball trials (p < 0.0035). To test the associations between pupil measurements and the triple-network model (SN, salience network; DAN, dorsal attention network; DMN, default mode network), we computed EC across nodes of these networks. (B) The oddball-modulated positive EC strength from SN to DAN correlated with higher TEPR of oddball trials (p < 0.0013). (C) The oddball-modulated negative EC strength from SN to DMN correlated with higher TEPR of oddball trials (p < 0.006).

### Involvement of Locus Coeruleus in Salience Processing

Given substantial evidence that pupil diameter is tightly coupled to the neuronal activity in the LC [3, 47], pupil diameter has been used as an index of the LC activity [35]. With the observed correlation between late-to-early network feedback signal and TEPR (Fig. 5A), we therefore hypothesized that LC might play a role in the interactions between large-scale cortical networks through the cortico-subcortical coupling. To test the involvement of LC in salience processing, we first examined the functional connectivity between LC and salience processing nodes. The LC showed significant functional connectivity with lSPL, lS1 (lSPL: t = 2.64, p = 0.017; lS1: t = 3.80, p = 0.001; nodes of early-time network and DAN) and mPFC-SMA (t = 3.15, p = 0.006; node of late-time network and SN), however, there are no significant results between LC and the other nodes (SI Results). With the functional coupling to the nodes of both early-time and late-time networks, this result indicates a mediator role of the LC that contributes to the causal interactions between these two networks. This result is also consistent with the observation that lSPL and mPFC-SMA are the hubs in the modulated EC, rendering their importance in salience processing.

Following these results on the involvement of LC with the late-to-early network feedback signal and the nodes of both DAN and SN, the final question we asked was whether this feedback signal reflects the interaction between DAN and SN, and whether LC mediates this interaction between the large-scale cortical networks. Given the vast amount of literature showing the close relationship between DAN, SN and default mode network (DMN), we fit the EEG data with the state-space model including the nodes of SN, DMN, and DAN, defined by the HCP-MMP (Human Connectome Project Multi-Modal Parcellation) atlas [36] (see SI Methods for details; EC results in Fig. S4). As expected, we found a significant positive correlation between the salience stimuli modulated SN-to-DAN positive network EC strength and TEPR (r = 0.6804, p = 0.0013; Fig. 5B; after controlling RT and ELBO: r = 0.5949, p = 0.0072). Interestingly, we also observed a significant negative correlation between the salience stimuli modulated SN-to-DMN negative network EC strength and TEPR (r = −0.6055, p = 0.006; Fig. 5C; after controlling RT and ELBO: r = −0.6820, p = 0.0013). These results are in line with the previous studies on the function of SN for the switching between anticorrelated networks [64, 97, 111]. In summary, the results indicate that the LC is involved in the switching between networks.

## Discussion

### Spatiotemporal Brain Networks in Salience Processing

We used EEG-informed fMRI analysis to map the spatiotemporal dynamics of neural substrates in salience processing. Specifically, the STV temporal information in the EEG was extracted at different time windows spanning the trial to explain the variance in the fMRI signal. This approach has been widely used to study a broad range of cognitive functions and human behaviors [66, 72, 107]. Compared to conventional fMRI analyses, this method allows us to temporally dissociate the stimuli-evoked brain activation, or even identify regions absent in the conventional analyses (canceled out due to a temporal integration effect) [37, 66, 72]. This work extends this approach by introducing the functional connectome into the framework for mapping the underlying spatiotemporal network organizations of these neural substrates. Functional connectomes have been used to elucidate intrinsic brain organizations [34], map lesion-associated networks [9, 33], model cognitive task activation [19, 45]. For example, seed-based functional connectivity was used in the lesion network localization to identify the specific symptom-associated brain network underlying spatially heterogeneous lesions with the same symptom [26]. In this study, based on the STV EEG-informed fMRI analysis and functional connectome network localization, we observed a spatiotemporal intrinsic network organization of the neural substrates in salience processing (Fig. 6). The involvement of these nodes and networks in salience processing or attention is consistent with prior studies: DAN, motor network, ECN and SN [49] and visual network [37].

**Figure 6.**
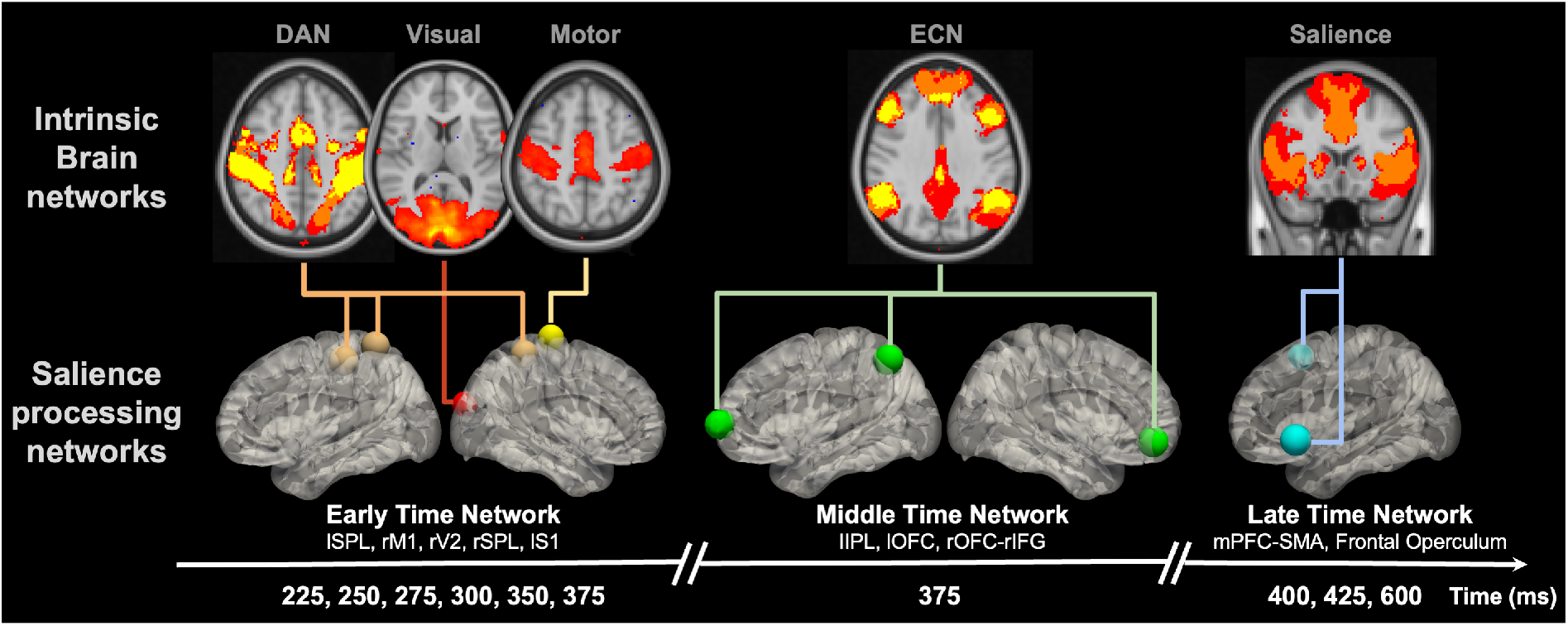
Neural cascades of salience processing and the spatiotemporal network organizations of salience processing nodes. Previous seed-based and node-by-node functional connectivity results suggest both spatial and temporal network organizations of the identified salience processing nodes, respectively. We hypothesized that the node-specific involvement of these functional networks might indicate a crucial role of these nodes in the temporally evolved processes of salience signal and the relationships between these networks. ECN, executive control network.

Given these observations, we inferred that the brain flexibly recruited specific nodes of distinct networks at different time windows spanning the trial according to the demand of specific cognitive processes and behavior responses. In the early-time windows, we observed a sustained activation of DAN subsystem from 225 ms to 375 ms, along with the rM1 at 225 ms and rV2 at 275 ms. Literatures have shown that DAN is associated with goal-driven attention and the role of linking them to appropriate motor responses [22]. The observed activation of the early-time network might reflect the interactions between these systems in salience processing. For example, the coactivation of rM1 and lSPL at 225 ms might reflect the interaction between DAN and the motor network and its role in linking stimuli and responses. Based on the previous evidence that DAN exhibits top-down influences on the sensory cortex [12, 22], we hypothesized that the involvement of the visual network might indicate the modulation of attentional resources distribution [86]. This hypothesis was also supported by the observed self-inhibition EC in rV2 (Fig. 4B). Similar to the reported temporal components underlying a visual spatial attention task in a MEG study [87], we also observed the involvement of ECN (375 ms) right after the early activation of parietal and visual areas. This node-specific involvement might indicate a crucial role of these nodes in the network and salience processing. For example, prior studies suggest the existence of common nodes (left orbital frontoinsula, mPFC, and right dorsolateral prefrontal cortex) between the ECN and SN [83]. Thus, the observed involvement of the left frontal operculum and mPFC-SMA in the late time might reflect their role in the interaction between ECN and SN, which facilitates the temporal transition from ECN to SN in the late time. Similarly, the coactivation of lS1 as a positive cluster and ECN nodes (lIPL, lOFC, and rOFC-rIFC) as negative clusters at 375 ms might indicate their role in the interaction between DAN and ECN. Future studies are needed to investigate the specific functions of these nodes in the interaction between brain networks.

### Cognitive Control in Salience Processing

Based on the anatomical locations of the nodes in the network, ECN and SN have also been named Lateral Frontoparietal Network (L-FPN) and Midcingulo-Insular Network (M-CIN), respectively [99]. Previous studies have shown that the ECN/L-FPN and SN/M-CIN are two prominent cognitive control networks, supporting the goal-directed cognition and behavior [20, 29, 30]. In these studies, L-FPN and M-CIN were named as frontoparietal network and cingulo-opercular network. To keep the terminology of the brain networks consistent in this study, we referred to ECN/L-FPN and SN/M-CIN following the guidelines in [99]. SN (cognitive domain name) and M-CIN (anatomical name) contains these core regions: bilateral anterior insula and anterior midcingulate cortex. ECN (cognitive domain name) and L-FPN (anatomical name) contains these core regions: lateral prefrontal cortex, anterior inferior parietal lobule, and intraparietal sulcus. ECN/L-FPN and SN/M-CIN are coactivated together as the multiple-demand system [25, 31] (sometimes also described as task-activation ensemble [83]) in goal-directed behaviors and tasks, such as working memory [52], cognitive control [21], and attention [108] tasks. However, converging evidence from the literature indicates distinct roles of ECN/L-FPN and SN/M-CIN in goal-directed behaviors. ECN/L-FPN acts as a flexible coordinator of goal-relevant information [18], and may underlie phasic control such as initiates exogenously triggered control, adaptive adjustments and executive functions [29, 79]. Whereas, SN/M-CIN is related to stable maintenance of task-set [29, 30] and tonic alertness [76], and has also been proposed a role in lending processing resources to help other goal-relevant networks [18].

In the present study, by leveraging STV temporal information in the EEG to tease apart the temporal neural processes in the goal-directed salience processing, we dissociated the stimuli-evoked coactivation of ECN/L-FPN and SN/M-CIN into two distinct temporal components (subsystem of ECN as the middle-time network activated at 375 ms, subsystem of SN as the late-time network activated during 400-600 ms). Along with this temporal dissociation, ECN/L-FPN and SN/M-CIN seemed to act with distinct roles in the salience processing, where ECN/L-FPN was activated preceding the behavior response (group-averaged median RT: 404 ms) and the activation of SN/M-CIN. This is consistent with the distinct functions of ECN/L-FPN and SN/M-CIN described in the literature. ECN/L-FPN acted as a phasic control (activated only at 375 ms), which provided the rapid control initiation to SN/M-CIN. This initiation might be mostly driven by the EC from lOFC to mPFC-SMA, as indicated by the strong connection strength (Fig. 4A). Whereas, SN/M-CIN was activated at multiple time points (400, 425, 600 ms) right after the response, indicating a stable maintenance of tonic alertness, which facilitates the better detection performance in the next coming trial [76, 77]. We propose that the involvement of SN in the late time windows might allow the brain to disengage the current trial and maintain the preparedness for the next upcoming stimulus. As proposed in a single-unit recording study, pre-SMA (mPFC-SMA) is associated with task switching by first suppressing irrelevant task-set and then boosting a controlled response with the relevant task-set [44, 80]. In the present study, we found mPFC-SMA deactivated starting at 400 ms (right after the response), which might reflect a suppression or inhibition of the current trial encoded task-set. With the proposed role of pre-SMA (mPFC-SMA) in conflict monitoring [10, 44], in the changing environment after the response, its involvement allows the brain to resolve the conflict between the current trial encoded task-set and the new environment, which faciliates task disengagement. This task-set suppressing and boosting role of the SN is consistent with the network switching theory [64]. The present findings are also in line with the proposed ‘‘windshield wiper” mechanism [76, 78] of the SN. In the anticipation of upcoming inputs, SN may employ such a mechanism to increase preparedness by clearing currently ongoing activity in multiple cortical areas. This was supported by the previously reported close relationship between SN activity, global α-oscillation power and the stable maintenance of tonic alertness [76, 78, 79]. As for the relationship between ECN/L-FPN and SN/M-CIN, the oddball-modulated EC results showed a stronger connectivity strength from ECN/L-FPN to SN/M-CIN (mid-to-late; positive: 0.2523; negative: 0.3145) compared to the connection from SN/M-CIN to ECN/L-FPN (late-to-mid; positive: 0.0165; negative: 0.1126). These results are consistent with the interactive dual-networks model in [28, 29], however an EC analysis with all the nodes of ECN/L-FPN and SN/M-CIN are needed to study the interactions between them. The present findings provided more evidence for the functions, relationship, and timescales of these two cognitive control networks (i.e. ECN/L-FPN and SN/M-CIN).

### Linking Networks Effective Connectivity and Pupillary Response: LC Mediates Network Reset

The LC-NE system has been proposed to modulate neural gain, attention and arousal [3]. Pupil diameter fluctuations, as a proxy of LC activity [47, 67], have been used to investigate how the ascending neuromodulator from the LC-NE system influences the cortex [68]. There is growing research on the relationship between brain measurements and pupil diameter, with evidence suggesting that pupil diameter fluctuations are associated with cortical membrane potential activity [61], EEG P300 component of the ERP [68], fMRI BOLD signal in the DMN, SN, thalamus, frontoparietal, visual and sensorimotor regions [81, 109], overall functional connectivity strength during exploration [91], global fluctuations in network structure [32], and global integration connectivity within a network of frontoparietal, striatal, and thalamic regions [85]. These findings shed light on the understanding of brain-pupil relationships and cortico-subcortical interactions, however, the role of the LC-NE system in the connectivity and interaction between specific brain networks within the context of a goal-driven task (e.g. salience processing) is less well understood. Here, by leveraging the high temporal resolution of the EEG, we used a state-space model for inferring the EC between brain networks involved in salience processing. We observed a very strong relationship between the late-to-early positive network interaction strength and TEPR (Fig. 5A). In the oddball paradigm with motor response, studies have shown that TEPR reflects not only bottom-up mechanisms but also top-down higher-order processing [46, 49]. Given the directionality of this TEPR-related network interaction, we propose that the phasic LC activity (indexed by TEPR) is associated with a feedback (top-down) signal from the late-time network (nodes of SN) to the early-time network (nodes of DAN, visual, and sensory motor network). Besides the close relationship between the LC activity and pupil diameter fluctuations, SN areas also exhibited close links to the LC-NE system and pupil measurements. The LC-NE system has shown to receive projections from the anterior cingulate cortex (ACC) and anterior insula (AI) [3, 22], and has robust functional connectivity with these SN nodes [55, 57]. The SN and pupil measurements have both been associated with task demands, efforts, difficulty [5, 71, 104], uncertainty and surprise [46], conflict and error processing [23], and anxiety [13, 83]. The involvement of SN, as a subsystem of the ventral attention network (VAN) [92, 99], aligns with evidence showing the involvement of VAN in both bottom-up stimulus saliency and top-down internal goals [22, 56, 106]. The observed feedback signal from SN may explain its proposed role in the global α-oscillation activity [76], which has been associated with top-down processes [4, 8, 41, 50, 70]. However, it warrants future studies to examine their relationship.

Even with widespread projections of LC neurons throughout the cortex, recent studies suggested that there are substantial specificity and heterogeneity in the projections [91, 93, 102]. For example, regions in DAN receive dense LC-NE inputs [7]. Our functional connectivity analysis suggests that the LC-NE system, with connections to both DAN and SN nodes, may act as a mediator in the top-down control from SN to DAN along with other early-time network nodes. Our results strongly support the network-reset theory, which proposes that the VAN (SN as a subsystem) marks behavior transitions and facilitates a network reset signal along with the phasic LC-NE activity (indexed by TEPR), to reconfigurate the DAN (part of the early-time network) for settling into another state in the new environment situation [11, 22, 27]. In light of recent studies in the LC-NE system effects on brain network reconfiguration [39, 110], more studies are needed to investigate the role of LC-NE system in the interaction between brain networks [73, 102].

### SN, DMN, and DAN in Salience Processing: LC Plays a Role in Network Switching

The anticorrelation between the DAN and the DMN has been characterized as a vital aspect of the human brain functional organization and dynamics [34], with DAN and DMN controlling environmentally directed and internally directed cognitive processes, respectively [22, 34]. Converging evidence suggests that the nodes of the SN are at the apex of the cortical hierarchy between these two anticorrelated networks [111], with a causal role in the dynamic switching between them [38, 90]. A triple network model has been proposed for these three core neurocognitive networks [62], serving as a networks framework for understanding psychopathology [62], cognitive aging [95], conscious perception [43]. However, the neural mechanisms underlying dynamic switching, and how can the SN have such a wide spread access to DAN and DMN for coordinating the switching between them, are not well understood. It has been proposed that the specialized von Economo neurons (VENs), which are only found in the SN, might relay the information from SN to other parts of the cortex [97]. However, the projections of VENs remain unknown, with some evidence suggesting projections to brainstem nuclei [17]. Hence, along with the close relationship between the SN and the LC-NE system discussed in the previous section, it is plausible to hypothesize that brainstem nuclei, such as the LC, may play a role in the dynamic switching of large-scale brain networks through the release of neuromodulatory neurotransmitters. Neuromodulation models of the LC have been proposed. In the adaptive-gain theory [3, 35], the LC receives task utility information from the ACC (SN node) and OFC, producing NE release at cortical target sites and adjust the gain. The network glutamate amplification of noradrenaline (GANE) model [58, 73] proposed that the SN recruits LC firing to enable NE local concentration modulation, accompanied by in parallel enhancement and suppression of large-scale brain networks. In support of our hypothesis, i.e. the LC facilitates network switching, we found that increased TEPR (index of phasic LC activity) is associated with a stronger positive EC from the SN to the DAN, and a stronger negative EC from the SN to the DMN (Fig. 5). This result confirmed the previous findings that the SN initiates the dynamic switching, and to our knowledge, this is the first study to show the role of the LC-NE system in the dynamic switching between anticorrelated networks. This suggests an cortico-subcortical integrated network switching (CS-INS) model, involving both the SN and the LC-NE system in the dynamic switching between the DAN and DMN (Fig. 7). This network switching model supports the notion of ‘enhancement’ and ‘suppression’ in the GANE model of the LC-NE system effects.

**Figure 7.**
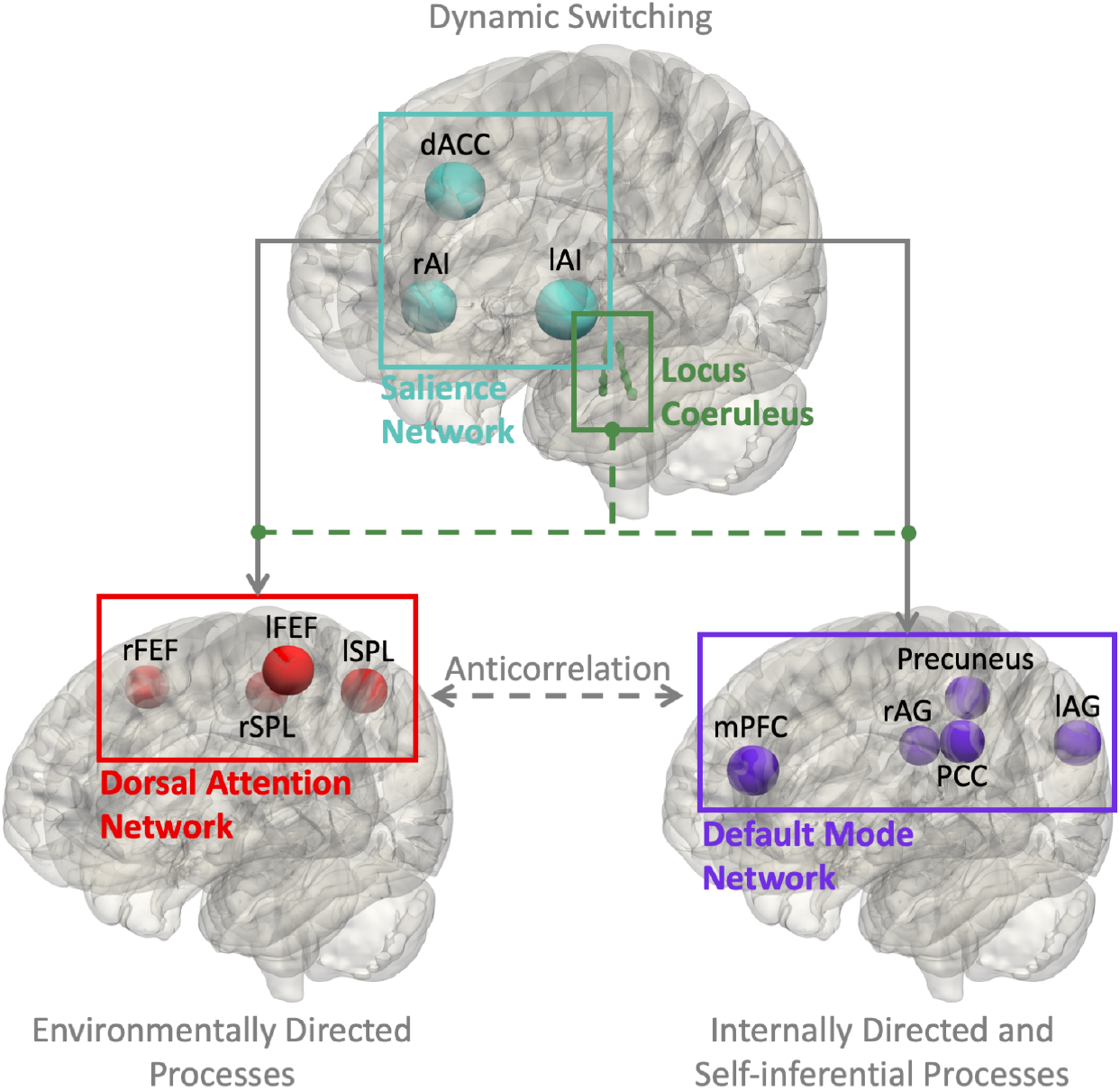
Cortico-subcortical integrated network switching (CS-INS) model. Previous brain-pupil relationships results aligned with the network switching model of SN in the literature, and also showed the role of the LC-NE system in the dynamic switching between anticorrelated networks (DAN and DMN). We hypothesized that the switching might be modulated by the release of the NE, as an effect of the ascending neuromodulation, which indicates that the SN and LC-NE system might cooperate and share an integrated role in salience processing. LC-NE, locus coeruleus-norepinephrine.

In support of the claims made previously, the strong relationship between the SN-to-DAN EC and TEPR suggests the role of the LC-NE system in mediating this network reset (or circuit-breaker) top-down control signal. Based on the involvement of the SN nodes as the late-time salience processing network (Fig. 6), and its higher causal hierarchy among the DMN and DAN [111], the CS-INS is thought to be involved in the late phases of the salience processing, reflecting a mark of behavior transition, task disengagement, and preparedness, initiated by the deactivation of SN. However, our findings do not rule out the possibility that the CS-INS system might also be involved at the other phases of the trial. For example, a recent study has found that the DAN, SN, visual and frontoparietal regions are involved in the early phase of the re-orienting, which has been interpreted as a network reset signal [89]. Different from our findings, these results might reflect the reorienting modulation in task engagement or simply bottom-up stimuli processing. Consistent with our hypothesis, the LC activation has been shown to be more closely aligned with the behavioral response than the stimulus onset [3, 73], and our previous work has also demonstrated that the DMN is involved in the late phase of the trial [107]. The DMN has been implicated in future planning [14], task switching [24], attention shifts [2], as well as other functions [14], and the LC-NE system has been proposed to modulate the DMN as a neural modulator of mind wandering [65]. Our results are consistent with hypothesis that the LC-NE system modulates the connectivity between the DMN and DAN [94], and also the proposed role as a ‘master switch’ [75]. Potential candidate mechanisms of the LC-NE modulation on the DMN and DAN are: 1) The LC-NE system modulates them through the heterogeneous spatial distribution of NE receptor types and densities across the cortex [101, 103]; 2) The heterogeneity in LC cell populations might be responsible for the targeted modulations of specific cortex areas [82, 93], for example, a modular organization in the LC with distinct efferent neural projection patterns has been reported [73, 100]; 3) The LC-NE system might interact with other subcortical nuclei in the modulation [102], such as thalamus, which has been associated with the LC activity [55, 57], SN activity [83], and the modulation of cortical networks connectivity [15, 40, 43, 69]. The work may inform future studies of the neural mechanisms underlying the LC-NE modulation on the cortex.

The CS-INS network model proposed here bridges the gap between the network models of the SN and the LC-NE system, which could potentially serve as a network paradigm for understanding the neural dynamics underlying cognitive functions, such as salience processing. With the neuronal basis of fast control in the VENs of SN [64] and the spatially diffuse projections from the LC [3], the CS-INS system is ideally suited to a variety of complex cognitive processes. Although our results shed light on the relationship between the LC-NE system and the cortical network switching in the context of an auditory oddball task, we speculate that the CS-INS system might play a more general role in cognitive functions, such as adaptations in environmental volatility [1, 13], brain state switches/variations [60, 61], cognitive control [18, 32, 35, 51]. Further work is needed to uncover the implications of the CS-INS model in various cognitive processes and neurological diseases. For example, previous studies have associated the activity in the DMN and DAN regions with exploitation and exploration, respectively [16, 53]. It has been proposed that the SN and the LC-NE system may play a role in the switching from exploitation to exploration [3, 16], hence, it will be interesting to test the CS-INS model in the exploration and exploitation tasks.

### Broader Implications of a Simultaneous Pupillometry-EEG-fMRI Study

In this study, we deployed a framework involving simultaneous recordings of pupillometry, EEG, and fMRI to investigate neural processes and interactions in salience processing. The high spatial resolution of fMRI data and functional connectivity analysis were utilized to map the neural substrates and the functional organizations. The EEG data with high temporal resolution, the single-trial analysis, and the state-space model were used to temporally ‘tag’ the neural substrates and infer the causal interactions. As a proxy of the LC activity, pupillometry was included to study the cortico-subcortical interactions. The methodological approaches and the CS-INS network model proposed here might promote further investigations on the brain dynamics underlying various cognitive processes and neurological diseases. For example, the role of the LC in cognitive control is intriguing but has not been fully explored. This might be due to the gap in current knowledge between the network models of the cortex and the models of the LC, such as how the LC interacts with the SN. Besides their critical contribution to attentional processing as demonstrated in this study, the CS-INS network model proposed here, or other cortico-subcortical network models, is also critical in the understanding of neurological diseases, such as Alzheimer’s disease (AD). As the first brain region in which AD-related pathology appears, the LC has been associated with cognitive decline and aging [59]. A recent study showed that the LC in the older population has reduced interaction with the SN, suggesting subsequent impairment in the initiation of network switching, and inferior ability in prioritizing the importance of incoming events [54].

### Limitations

Making causal inferences from correlation is challenging [74]. We used EC to infer the causal couplings between brain networks, yet the directionality in the cortico-subcortical interactions between the LC and cortex regions remain unclear. Based on the involvement of the SN in the late time of the trial observed in our data, we hypothesized that the network switching might be modulated by the release of the NE, as an effect of the ascending neuromodulation. Whereas, in the adaptive gain theory, the LC is proposed to receive inputs from ACC and OFC, with the release of NE at cortical target sites [3]. Further investigations will be needed to better understand the directionality in the interactions between the LC and cortex regions.

Pupillometry has long been used to index the LC activity in previous studies [3, 35, 46, 68] and also in the current study, though it is challenging, as other neural circuits are involved in controlling pupil diameter as well [46]. For example, shifts of attention are mediated in part by the superior colliculus (SC). In the previous studies, the LC rather than the SC showed neural spiking responses to unexpected auditory events [47]. In an oddball task fMRI study, the pupil size fluctuations have been associated with the BOLD activity in the LC [67]. In this study, we utilized this well-studied oddball paradigm, where the coupling between pupillometry and the LC activity has been shown, to investigate the pupil-indexed activity in the LC. However, this does not rule out the possibility that other subcortical nuclei, such as thalamus, or other neuromodulators, such as acetylcholine, might contribute to pupil diameter fluctuations or interact with the LC-NE system. Future studies are needed for direct neuroimaging of the LC, however, it is challenging due to the excessive physiological noise and distortion in brainstem imaging [6], and the difficulty in the localization of the LC [57].

In this study, we made inferences on the role of the pupil-indexed LC activity in salience processing, based on their interaction with the task-related neural substrates. However, besides the attentional processing of salient stimuli, pupil-indexed LC activity has also been associated to changes in arousal. Though these two LC associated processes, i.e. attention and arousal, have been shown to be independent [105], an important future direction will be accounting for the LC associated arousal, and assessing its relationship to the cortex. In future studies, novel tasks can be devised to dissociate pupil-indexed LC activity in attention and arousal.

## Materials and Methods

Nineteen subjects (mean age ± SD = 25.9 ± 3.6 years, female/male = 13/6; SD, standard deviation) were included in all the analyses. The experimental design of our study and the recruitment process were approved by Columbia University institutional review board. An auditory oddball paradigm with 80% standard and 20% oddball (target) stimuli was performed, where standard stimuli were pure tones with a frequency of 350 Hz, and the oddball stimuli were broadband (laser gun) sounds. A 3T Siemens Prisma scanner was used to acquire pupillometry, EEG and fMRI with a 64 channel head coil. Details of experimental procedures, data preprocessing and data analysis are provided in SI Methods.

## Acknowledgments

This work was supported by an Army Research Laboratory Cooperative Agreement W911NF-10-2-0022 and a Vannevar Bush Faculty Fellowship from the US Department of Defense (N00014-20-1-2027).

## References

1. J. Y. Angela and P. Dayan. Uncertainty, neuromodulation, and attention. Neuron, 46(4):681–692, 2005.

2. J. T. Arsenault, N. Caspari, R. Vandenberghe, and W. Vanduffel. Attention shifts recruit the monkey default mode network. Journal of Neuroscience, 38(5):1202–1217, 2018.

3. G. Aston-Jones and J. D. Cohen. An integrative theory of locus coeruleus-norepinephrine function: adaptive gain and optimal performance. Annu. Rev. Neurosci., 28:403–450, 2005.

4. A. M. Bastos, W. M. Usrey, R. A. Adams, G. R. Mangun, P. Fries, and K. J. Friston. Canonical microcircuits for predictive coding. Neuron, 76(4):695–711, 2012.

5. J. Beatty. Task-evoked pupillary responses, processing load, and the structure of processing resources. Psychological Bulletin, 91(2):276, 1982.

6. F. Beissner. Functional MRI of the brainstem: common problems and their solutions. Clinical Neuroradiology, 25(2):251–257, 2015.

7. E. E. Benarroch. The locus ceruleus norepinephrine system: functional organization and potential clinical significance. Neurology, 73(20):1699–1704, 2009.

8. M. Benedek, S. Bergner, T. Könen, A. Fink, and A. C. Neubauer. EEG alpha synchronization is related to top-down processing in convergent and divergent thinking. Neuropsychologia, 49(12):3505–3511, 2011.

9. A. D. Boes, S. Prasad, H. Liu, Q. Liu, A. Pascual-Leone, V. S. Caviness Jr, and M. D. Fox. Network localization of neurological symptoms from focal brain lesions. Brain, 138(10):3061–3075, 2015.

10. M. M. Botvinick. Conflict monitoring and decision making: reconciling two perspectives on anterior cingulate function. Cognitive, Affective, & Behavioral Neuroscience, 7(4):356–366, 2007.

11. S. Bouret and S. J. Sara. Network reset: a simplified overarching theory of locus coeruleus noradrenaline function. Trends in Neurosciences, 28(11):574–582, 2005.

12. S. L. Bressler, W. Tang, C. M. Sylvester, G. L. Shulman, and M. Corbetta. Top-down control of human visual cortex by frontal and parietal cortex in anticipatory visual spatial attention. Journal of Neuroscience, 28(40):10056–10061, 2008.

13. M. Browning, T. E. Behrens, G. Jocham, J. X. O’reilly, and S. J. Bishop. Anxious individuals have difficulty learning the causal statistics of aversive environments. Nature Neuroscience, 18(4):590–596, 2015.

14. R. L. Buckner, J. R. Andrews-Hanna, and D. L. Schacter. The brain’s default network: anatomy, function, and relevance to disease. 2008.

15. R. L. Buckner and L. M. DiNicola. The brain’s default network: updated anatomy, physiology and evolving insights. Nature Reviews Neuroscience, 20(10):593–608, 2019.

16. K. Chakroun, D. Mathar, A. Wiehler, F. Ganzer, and J. Peters. Dopaminergic modulation of the exploration/exploitation trade-off in human decision-making. eLife, 9:e51260, 2020.

17. I. Cobos and W. W. Seeley. Human von Economo neurons express transcription factors associated with layer V subcerebral projection neurons. Cerebral Cortex, 25(1):213–220, 2015.

18. C. V. Cocuzza, T. Ito, D. Schultz, D. S. Bassett, and M. W. Cole. Flexible coordinator and switcher hubs for adaptive task control. Journal of Neuroscience, 40(36):6949–6968, 2020.

19. M. W. Cole, T. Ito, D. S. Bassett, and D. H. Schultz. Activity flow over resting-state networks shapes cognitive task activations. Nature Neuroscience, 19(12):1718–1726, 2016.

20. M. W. Cole, J. R. Reynolds, J. D. Power, G. Repovs, A. Anticevic, and T. S. Braver. Multi-task connectivity reveals flexible hubs for adaptive task control. Nature Neuroscience, 16(9):1348–1355, 2013.

21. M. W. Cole and W. Schneider. The cognitive control network: Integrated cortical regions with dissociable functions. NeuroImage, 37(1):343–360, 2007.

22. M. Corbetta, G. Patel, and G. L. Shulman. The reorienting system of the human brain: from environment to theory of mind. Neuron, 58(3):306–324, 2008.

23. H. D. Critchley, J. Tang, D. Glaser, B. Butterworth, and R. J. Dolan. Anterior cingulate activity during error and autonomic response. NeuroImage, 27(4):885–895, 2005.

24. B. M. Crittenden, D. J. Mitchell, and J. Duncan. Recruitment of the default mode network during a demanding act of executive control. eLife, 4:e06481, 2015.

25. B. M. Crittenden, D. J. Mitchell, and J. Duncan. Task encoding across the multiple demand cortex is consistent with a frontoparietal and cingulo-opercular dual networks distinction. Journal of Neuroscience, 36(23):6147–6155, 2016.

26. R. R. Darby, A. Horn, F. Cushman, and M. D. Fox. Lesion network localization of criminal behavior. Proceedings of the National Academy of Sciences, 115(3):601–606, 2018.

27. P. Dayan and A. J. Yu. Phasic norepinephrine: a neural interrupt signal for unexpected events. Network: Computation in Neural Systems, 17(4):335–350, 2006.

28. N. U. Dosenbach, D. A. Fair, A. L. Cohen, B. L. Schlaggar, and S. E. Petersen. A dual-networks architecture of top-down control. Trends in Cognitive Sciences, 12(3):99–105, 2008.

29. N. U. Dosenbach, D. A. Fair, F. M. Miezin, A. L. Cohen, K. K. Wenger, R. A. Dosenbach, M. D. Fox, A. Z. Snyder, J. L. Vincent, M. E. Raichle, et al. Distinct brain networks for adaptive and stable task control in humans. Proceedings of the National Academy of Sciences, 104(26):11073–11078, 2007.

30. N. U. Dosenbach, K. M. Visscher, E. D. Palmer, F. M. Miezin, K. K. Wenger, H. C. Kang, E. D. Burgund, A. L. Grimes, B. L. Schlaggar, and S. E. Petersen. A core system for the implementation of task sets. Neuron, 50(5):799–812, 2006.

31. J. Duncan. The multiple-demand (MD) system of the primate brain: mental programs for intelligent behaviour. Trends in Cognitive Sciences, 14(4):172–179, 2010.

32. E. Eldar, J. D. Cohen, and Y. Niv. The effects of neural gain on attention and learning. Nature Neuroscience, 16(8):1146–1153, 2013.

33. M. D. Fox. Mapping symptoms to brain networks with the human connectome. New England Journal of Medicine, 379(23):2237–2245, 2018.

34. M. D. Fox, A. Z. Snyder, J. L. Vincent, M. Corbetta, D. C. Van Essen, and M. E. Raichle. The human brain is intrinsically organized into dynamic, anticorrelated functional networks. Proceedings of the National Academy of Sciences, 102(27):9673–9678, 2005.

35. M. S. Gilzenrat, S. Nieuwenhuis, M. Jepma, and J. D. Cohen. Pupil diameter tracks changes in control state predicted by the adaptive gain theory of locus coeruleus function. Cognitive, Affective, & Behavioral Neuroscience, 10(2):252–269, 2010.

36. M. F. Glasser, T. S. Coalson, E. C. Robinson, C. D. Hacker, J. Harwell, E. Yacoub, K. Ugurbil, J. Andersson, C. F. Beckmann, M. Jenkinson, et al. A multi-modal parcellation of human cerebral cortex. Nature, 536(7615):171–178, 2016.

37. R. I. Goldman, C.-Y. Wei, M. G. Philiastides, A. D. Gerson, D. Friedman, T. R. Brown, and P. Sajda. Single-trial discrimination for integrating simultaneous EEG and fMRI: identifying cortical areas contributing to trial-to-trial variability in the auditory oddball task. NeuroImage, 47(1):136–147, 2009.

38. N. Goulden, A. Khusnulina, N. J. Davis, R. M. Bracewell, A. L. Bokde, J. P. McNulty, and P. G. Mullins. The salience network is responsible for switching between the default mode network and the central executive network: replication from DCM. NeuroImage, 99:180–190, 2014.

39. C. Guedj, E. Monfardini, A. J. Reynaud, A. Farnè, M. Meunier, and F. Hadj-Bouziane. Boosting norepinephrine transmission triggers flexible reconfiguration of brain networks at rest. Cerebral Cortex, 27(10):4691–4700, 2017.

40. M. M. Halassa and S. Kastner. Thalamic functions in distributed cognitive control. Nature Neuroscience, 20(12):1669–1679, 2017.

41. M. Halgren, I. Ulbert, H. Bastuji, D. Fabó, L. Erőss, M. Rey, O. Devinsky, W. K. Doyle, R. Mak-McCully, E. Halgren, et al. The generation and propagation of the human alpha rhythm. Proceedings of the National Academy of Sciences, 116(47):23772–23782, 2019.

42. H. A. Harsay, M. Spaan, J. G. Wijnen, and K. R. Ridderinkhof. Error awareness and salience processing in the oddball task: shared neural mechanisms. Frontiers in Human Neuroscience, 6:246, 2012.

43. Z. Huang, V. Tarnal, P. E. Vlisides, E. L. Janke, A. M. McKinney, P. Picton, G. A. Mashour, and A. G. Hudetz. Anterior insula regulates brain network transitions that gate conscious access. Cell Reports, 35(5):109081, 2021.

44. M. Isoda and O. Hikosaka. Switching from automatic to controlled action by monkey medial frontal cortex. Nature Neuroscience, 10(2):240–248, 2007.

45. T. Ito, K. R. Kulkarni, D. H. Schultz, R. D. Mill, R. H. Chen, L. I. Solomyak, and M. W. Cole. Cognitive task information is transferred between brain regions via resting-state network topology. Nature Communications, 8(1):1–14, 2017.

46. S. Joshi and J. I. Gold. Pupil size as a window on neural substrates of cognition. Trends in Cognitive Sciences, 24(6):466–480, 2020.

47. S. Joshi, Y. Li, R. M. Kalwani, and J. I. Gold. Relationships between pupil diameter and neuronal activity in the locus coeruleus, colliculi, and cingulate cortex. Neuron, 89(1):221–234, 2016.

48. C. Justen and C. Herbert. The spatio-temporal dynamics of deviance and target detection in the passive and active auditory oddball paradigm: a sLORETA study. BMC Neuroscience, 19(1):1–18, 2018.

49. H. Kim. Involvement of the dorsal and ventral attention networks in oddball stimulus processing: A meta-analysis. Human Brain Mapping, 35(5):2265–2284, 2014.

50. W. Klimesch. Alpha-band oscillations, attention, and controlled access to stored information. Trends in Cognitive Sciences, 16(12):606–617, 2012.

51. S. Köhler, K.-J. Bär, and G. Wagner. Differential involvement of brainstem noradrenergic and midbrain dopaminergic nuclei in cognitive control. Human Brain Mapping, 37(6):2305–2318, 2016.

52. B. Krasnow, L. Tamm, M. D. Greicius, T. Yang, G. H. Glover, A. L. Reiss, and V. Menon. Comparison of fMRI activation at 3 and 1.5 T during perceptual, cognitive, and affective processing. NeuroImage, 18(4):813–826, 2003.

53. D. Laureiro-Martínez, N. Canessa, S. Brusoni, M. Zollo, T. Hare, F. Alemanno, and S. F. Cappa. Frontopolar cortex and decision-making efficiency: comparing brain activity of experts with different professional background during an exploration-exploitation task. Frontiers in Human Neuroscience, 7:927, 2014.

54. T.-H. Lee, S. H. Kim, B. Katz, and M. Mather. The decline in intrinsic connectivity between the salience network and locus coeruleus in older adults: implications for distractibility. Frontiers in Aging Neuroscience, 12:2, 2020.

55. T. Liebe, J. Kaufmann, M. Li, M. Skalej, G. Wagner, and M. Walter. In vivo anatomical mapping of human locus coeruleus functional connectivity at 3 T MRI. Human Brain Mapping, 41(8):2136–2151, 2020.

56. N. M. Long and B. A. Kuhl. Bottom-up and top-down factors differentially influence stimulus representations across large-scale attentional networks. Journal of Neuroscience, 38(10):2495–2504, 2018.

57. V. Mäki-Marttunen and T. Espeseth. Uncovering the locus coeruleus: comparison of localization methods for functional analysis. NeuroImage, 224:117409, 2021.

58. M. Mather, D. Clewett, M. Sakaki, and C. W. Harley. Norepinephrine ignites local hotspots of neuronal excitation: How arousal amplifies selectivity in perception and memory. Behavioral and Brain Sciences, 39, 2016.

59. M. Mather and C. W. Harley. The locus coeruleus: essential for maintaining cognitive function and the aging brain. Trends in Cognitive Sciences, 20(3):214–226, 2016.

60. D. A. McCormick, D. B. Nestvogel, and B. J. He. Neuromodulation of brain state and behavior. Annual Review of Neuroscience, 43:391–415, 2020.

61. M. J. McGinley, S. V. David, and D. A. McCormick. Cortical membrane potential signature of optimal states for sensory signal detection. Neuron, 87(1):179–192, 2015.

62. V. Menon. Large-scale brain networks and psychopathology: a unifying triple network model. Trends in Cognitive Sciences, 15(10):483–506, 2011.

63. V. Menon. Salience network: Brain mapping: An encyclopedic reference, 2015.

64. V. Menon and L. Q. Uddin. Saliency, switching, attention and control: a network model of insula function. Brain Structure and Function, 214(5-6):655–667, 2010.

65. M. Mittner, G. E. Hawkins, W. Boekel, and B. U. Forstmann. A neural model of mind wandering. Trends in Cognitive Sciences, 20(8):570–578, 2016.

66. J. Muraskin, T. R. Brown, J. M. Walz, T. Tu, B. Conroy, R. I. Goldman, and P. Sajda. A multimodal encoding model applied to imaging decision-related neural cascades in the human brain. NeuroImage, 180:211–222, 2018.

67. P. R. Murphy, R. G. O’connell, M. O’sullivan, I. H. Robertson, and J. H. Balsters. Pupil diameter covaries with BOLD activity in human locus coeruleus. Human Brain Mapping, 35(8):4140–4154, 2014.

68. P. R. Murphy, I. H. Robertson, J. H. Balsters, and R. G. O’connell. Pupillometry and P3 index the locus coeruleus-noradrenergic arousal function in humans. Psychophysiology, 48(11):1532–1543, 2011.

69. M. Nakajima and M. M. Halassa. Thalamic control of functional cortical connectivity. Current Opinion in Neurobiology, 44:127–131, 2017.

70. S. Palva and J. M. Palva. Functional roles of alpha-band phase synchronization in local and large-scale cortical networks. Frontiers in Psychology, 2:204, 2011.

71. M. G. Philiastides and P. Sajda. EEG-informed fMRI reveals spatiotemporal characteristics of perceptual decision making. Journal of Neuroscience, 27(48):13082–13091, 2007.

72. M. G. Philiastides, T. Tu, and P. Sajda. Inferring macroscale brain dynamics via fusion of simultaneous EEG-fMRI. Annual Review of Neuroscience, 44, 2021.

73. G. R. Poe, S. Foote, O. Eschenko, J. P. Johansen, S. Bouret, G. Aston-Jones, C. W. Harley, D. Manahan-Vaughan, D. Weinshenker, R. Valentino, et al. Locus coeruleus: a new look at the blue spot. Nature Reviews Neuroscience, 21(11):644–659, 2020.

74. A. T. Reid, D. B. Headley, R. D. Mill, R. Sanchez-Romero, L. Q. Uddin, D. Marinazzo, D. J. Lurie, P. A. Valdés-Sosa, S. J. Hanson, B. B. Biswal, et al. Advancing functional connectivity research from association to causation. Nature Neuroscience, 22(11):1751–1760, 2019.

75. J. A. Ross and E. J. Van Bockstaele. The locus coeruleus-norepinephrine system in stress and arousal: unraveling historical, current, and future perspectives. Frontiers in Psychiatry, 11:1581, 2021.

76. S. Sadaghiani and M. D’Esposito. Functional characterization of the cingulo-opercular network in the maintenance of tonic alertness. Cerebral Cortex, 25(9):2763–2773, 2015.

77. S. Sadaghiani, G. Hesselmann, and A. Kleinschmidt. Distributed and antagonistic contributions of ongoing activity fluctuations to auditory stimulus detection. Journal of Neuroscience, 29(42):13410–13417, 2009.

78. S. Sadaghiani and A. Kleinschmidt. Brain networks and α-oscillations: structural and functional foundations of cognitive control. Trends in Cognitive Sciences, 20(11):805–817, 2016.

79. S. Sadaghiani, R. Scheeringa, K. Lehongre, B. Morillon, A.-L. Giraud, M. d’Esposito, and A. Kleinschmidt. Alpha-band phase synchrony is related to activity in the fronto-parietal adaptive control network. Journal of Neuroscience, 32(41):14305–14310, 2012.

80. K. Sakai. Task set and prefrontal cortex. Annu. Rev. Neurosci., 31:219–245, 2008.

81. M. Schneider, P. Hathway, L. Leuchs, P. G. Sämann, M. Czisch, and V. I. Spoormaker. Spontaneous pupil dilations during the resting state are associated with activation of the salience network. NeuroImage, 139:189–201, 2016.

82. L. A. Schwarz and L. Luo. Organization of the locus coeruleus-norepinephrine system. Current Biology, 25(21):R1051–R1056, 2015.

83. W. W. Seeley, V. Menon, A. F. Schatzberg, J. Keller, G. H. Glover, H. Kenna, A. L. Reiss, and M. D. Greicius. Dissociable intrinsic connectivity networks for salience processing and executive control. Journal of Neuroscience, 27(9):2349–2356, 2007.

84. J. M. Shine. Neuromodulatory influences on integration and segregation in the brain. Trends in Cognitive Sciences, 23(7):572–583, 2019.

85. J. M. Shine, P. G. Bissett, P. T. Bell, O. Koyejo, J. H. Balsters, K. J. Gorgolewski, C. A. Moodie, and R. A. Poldrack. The dynamics of functional brain networks: integrated network states during cognitive task performance. Neuron, 92(2):544–554, 2016.

86. S. Shomstein and S. Yantis. Control of attention shifts between vision and audition in human cortex. Journal of Neuroscience, 24(47):10702–10706, 2004.

87. G. V. Simpson, D. L. Weber, C. L. Dale, D. Pantazis, S. L. Bressler, R. M. Leahy, and T. L. Luks. Dynamic activation of frontal, parietal, and sensory regions underlying anticipatory visual spatial attention. Journal of Neuroscience, 31(39):13880–13889, 2011.

88. S. M. Smith, P. T. Fox, K. L. Miller, D. C. Glahn, P. M. Fox, C. E. Mackay, N. Filippini, K. E. Watkins, R. Toro, A. R. Laird, et al. Correspondence of the brain’s functional architecture during activation and rest. Proceedings of the National Academy of Sciences, 106(31):13040–13045, 2009.

89. S. Spadone, V. Betti, C. Sestieri, V. Pizzella, M. Corbetta, and S. Della Penna. Spectral signature of attentional reorienting in the human brain. NeuroImage, page 118616, 2021.

90. D. Sridharan, D. J. Levitin, and V. Menon. A critical role for the right fronto-insular cortex in switching between central-executive and default-mode networks. Proceedings of the National Academy of Sciences, 105(34):12569–12574, 2008.

91. N. Tardiff, J. D. Medaglia, D. S. Bassett, and S. L. Thompson-Schill. The modulation of brain network integration and arousal during exploration. NeuroImage, 240:118369, 2021.

92. B. Thomas Yeo, F. M. Krienen, J. Sepulcre, M. R. Sabuncu, D. Lashkari, M. Hollinshead, J. L. Roffman, J.W. Smoller, L. Zöllei, J. R. Polimeni, et al. The organization of the human cerebral cortex estimated by intrinsic functional connectivity. Journal of Neurophysiology, 106(3):1125–1165, 2011.

93. N. K. Totah, N. K. Logothetis, and O. Eschenko. Noradrenergic ensemble-based modulation of cognition over multiple timescales. Brain Research, 1709:50–66, 2019.

94. J. S. Tsukahara and R. W. Engle. Fluid intelligence and the locus coeruleus-norepinephrine system. Proceedings of the National Academy of Sciences, 118(46), 2021.

95. K. A. Tsvetanov, R. N. Henson, L. K. Tyler, A. Razi, L. Geerligs, T. E. Ham, J. B. Rowe, et al. Extrinsic and intrinsic brain network connectivity maintains cognition across the lifespan despite accelerated decay of regional brain activation. Journal of Neuroscience, 36(11):3115–3126, 2016.

96. T. Tu, J. Paisley, S. Haufe, and P. Sajda. A state-space model for inferring effective connectivity of latent neural dynamics from simultaneous EEG/fMRI. Advances in Neural Information Processing Systems, 32:4662–4671, 2019.

97. L. Q. Uddin. Salience processing and insular cortical function and dysfunction. Nature Reviews Neuroscience, 16(1):55–61, 2015.

98. L. Q. Uddin. Salience network of the human brain. Academic Press, 2016.

99. L. Q. Uddin, B. T. Yeo, and R. N. Spreng. Towards a universal taxonomy of macro-scale functional human brain networks. Brain Topography, 32(6):926–942, 2019.

100. A. Uematsu, B. Z. Tan, E. A. Ycu, J. S. Cuevas, J. Koivumaa, F. Junyent, E. J. Kremer, I. B. Witten, K. Deisseroth, and J. P. Johansen. Modular organization of the brainstem noradrenaline system coordinates opposing learning states. Nature Neuroscience, 20(11):1602–1611, 2017.

101. R. L. van den Brink, S. Nieuwenhuis, and T. H. Donner. Amplification and suppression of distinct brainwide activity patterns by catecholamines. Journal of Neuroscience, 38(34):7476–7491, 2018.

102. R. L. van den Brink, T. Pfeffer, and T. H. Donner. Brainstem modulation of large-scale intrinsic cortical activity correlations. Frontiers in Human Neuroscience, 13:340, 2019.

103. R. L. van den Brink, T. Pfeffer, C. M. Warren, P. R. Murphy, K.-D. Tona, N. J. van der Wee, E. Giltay, M. S. van Noorden, S. A. Rombouts, T. H. Donner, et al. Catecholaminergic neuromodulation shapes intrinsic MRI functional connectivity in the human brain. Journal of Neuroscience, 36(30):7865–7876, 2016.

104. E. Vassena, M. Silvetti, C. N. Boehler, E. Achten, W. Fias, and T. Verguts. Overlapping neural systems represent cognitive effort and reward anticipation. PLoS One, 9(3):e91008, 2014.

105. E. M. Vazey, D. E. Moorman, and G. Aston-Jones. Phasic locus coeruleus activity regulates cortical encoding of salience information. Proceedings of the National Academy of Sciences, 115(40):E9439–E9448, 2018.

106. S. Vossel, J. J. Geng, and G. R. Fink. Dorsal and ventral attention systems: distinct neural circuits but collaborative roles. The Neuroscientist, 20(2):150–159, 2014.

107. J. M. Walz, R. I. Goldman, M. Carapezza, J. Muraskin, T. R. Brown, and P. Sajda. Simultaneous EEG-fMRI reveals a temporal cascade of task-related and default-mode activations during a simple target detection task. NeuroImage, 102:229–239, 2014.

108. D. Weissman, L. Warner, and M. Woldorff. The neural mechanisms for minimizing cross-modal distraction. Journal of Neuroscience, 24(48):10941–10949, 2004.

109. D. Yellin, A. Berkovich-Ohana, and R. Malach. Coupling between pupil fluctuations and resting-state fMRI uncovers a slow build-up of antagonistic responses in the human cortex. NeuroImage, 106:414–427, 2015.

110. V. Zerbi, A. Floriou-Servou, M. Markicevic, Y. Vermeiren, O. Sturman, M. Privitera, L. von Ziegler, K. D. Ferrari, B. Weber, P. P. De Deyn, et al. Rapid reconfiguration of the functional connectome after chemogenetic locus coeruleus activation. Neuron, 103(4):702–718, 2019.

111. Y. Zhou, K. J. Friston, P. Zeidman, J. Chen, S. Li, and A. Razi. The hierarchical organization of the default, dorsal attention and salience networks in adolescents and young adults. Cerebral Cortex, 28(2):726–737, 2018.

