## Supplementary Information for "Spatiotemporal dissociation of brain network connectivity in salience processing: A simultaneous pupillometry-EEG-fMRI study"

### Methods

#### Participants and Experimental Paradigm

Twenty-five healthy subjects were recruited in this study and six of them were excluded from further analyses due to 1) missing neuroimaging data; 2) abnormality in the acquired neuroimaging data; 3) excessive movement; 4) inability to complete the task. Data from the remaining nineteen subjects (mean age $\pm$ SD = 25.9 $\pm$ 3.6 years, female/male = 13/6; SD, standard deviation) were included in the analyses. All subjects had normal or corrected-to-normal vision and no history of psychiatric illness or head injury. The experimental design of our study and the recruitment process were approved by Columbia University institutional review board.

An auditory oddball paradigm with 80% standard and 20% oddball (target) stimuli was performed, where standard stimuli were pure tones with a frequency of 350 Hz, and the oddball stimuli were broadband (laser gun) sounds. The inter-trial interval were in the range between 2 s and 3 s drawn from a uniform distribution, and each stimulus lasted for 200 ms. Subjects were first trained outside of the scanner to learn and perform the task comfortably and accurately on short training runs. All subjects performed the task correctly during training. During the data acquisition, stimuli were presented through MR compatible earphones, and subjects were instructed to maintain the fixation on the screen to a fixation target, and press a button (MR-compatible button box; PYKA, Current Designs, PA, USA) with their right index finger as soon as they heard the oddball sound. And subjects were instructed to ignore standard tones. All subjects responded to the task correctly with an accuracy of 99.4% $\pm$ 0.1% (mean $\pm$ SD) in detecting oddballs. Every subject was scheduled to complete five runs (105 trials per run), with an average of 4.7 runs per subject (range from three to five, SD = 0.7 runs) acquired in the experiment. The auditory oddball experimental task paradigm is illustrated in the first row of Fig. S5., where the first five trials were constrained to be standard stimuli, and no consecutive oddball trials was allowed.

#### Data Acquisition

A 3T Siemens Prisma scanner was used to acquire pupillometry, EEG and fMRI with a 64 channel head coil. Pupillometry was recorded with a MR-compatible EyeLink 1000 Plus in Long Range Mount, at a sampling rate of 1 kHz. EEG was recorded with a 64 channel BrainAmp MR Plus system (Brain Products, Germany), at a sampling rate of 5 kHz. The 64 channels include 63 cap electrodes and 1 ECG electrode in an extended 10-20 configuration with ground electrode at AFz and reference electrode at FCz. Functional MRI data were collected with T2*-weighted echo planar imaging (EPI) interleaved slice acquisition (TR = 2100 ms; TE = 25 ms; voxel size $3 \times 3 \times 3$mm; Matrix Size = $64 \times 64 \times 42$; 150 volumes). For localization and registration purposes, we collected T1-weighted structural image (MPRAGE, TR = 2300 ms; TE = 3.95 ms; voxel size = $1 \times 1 \times 1$ mm; Matrix Size = $176 \times 248 \times 256$) and T2*-weighted high-resolution EPI (TR = 6000 ms; TE = 30 ms; voxel size $2 \times 2 \times 3$ mm; Matrix Size = $96 \times 96 \times$

42; single-volume). For the localization of LC, we also collected neuromelanin-sensitive MRI data using T1-weighted turbo-spin-echo (TSE) acquisition (TR = 600ms; TE = 14ms; voxel size 0.43 × 0.43 × 6mm; Matrix Size = 416 × 512 × 5).

#### EEG Analysis

##### Data preprocessing

Visual inspection was first performed, to make sure the raw EEG signal was not contaminated by confounding factors, including TR volume jitter and data saturation. An average artifact template subtraction approach was then used to remove the gradient artifact with Brain Products' Analyzer2 data processing software [1,2]. The data were then down-sampled to 500 Hz. After the gradient artifact removal, a tenth order median filter was applied, to reject any residual gradient artifact. Then, the EEG data were filtered with a fourth order bandpass Butterworth filter (0.5 Hz to 50 Hz), to remove DC drift and high frequency noise. After filtering, each subject's EEG data were concatenated over runs for the application of ballistocardiogram artifact (BCG) removal. Specifically, QRS detection was first carried out, and then the BCG was removed with EEGLAB's FMRIB plugin (simple mean approach). The BCG removed data were then re-referenced to the common average. The final step of preprocessing is blink artifact removal, where independent component analysis (ICA) was performed using EEGLAB's ICA function to compute ICs, manually identify and remove the blink ICs [10]. The preprocessed EEG data were epoched identically as the pupillometry data from 500 ms before stimulus to 2000 ms following the stimulus. Then, baseline correction was carried out by removing the mean baseline value, which was computed from 500 ms before the stimulus onset to the stimulus onset for each epoch. We performed a trial rejection using a probability distribution based criteria. Specifically, we rejected trials where the EEG signal from a single channel is outside of 6SD, and the EEG signals from all channels are outside of 2SD. We also rejected trials where subjects incorrectly responded or failed to respond.

##### Single-trial analysis

A single-trial analysis with the sliding window approach was performed on the preprocessed EEG signal amplitude [14]. Specifically, with a linear classifier maximally discriminating the target versus standard trials, a hyper-plane was learnt to project the multidimensional EEG signal into lowdimensional EEG single-trial variability (STV) discriminating components. Given the EEG signal, $y_i(t)$ at time $t$, where $i = 1, 2 \ldots, T$ denotes trial index, logistic regression was used to learn the projection weights $w(\tau)$. The lowdimensional EEG STV discriminating components will be: $d_i(\tau) = \frac{1}{N} \sum_{t=\tau-\frac{N}{2}}^{\tau+\frac{N}{2}} w(\tau)^T y_i(t)$, where $N = 50ms$ denotes window width, the window center $\tau$ was shifted from 0 to 1000 ms with respect to the stimulus onset in 25 ms increments. For each temporal window, the classifier performance was assessed with the area under the receiver operating characteristic curve (AUC) using leave-one-out (LOO) cross-validation. A permutation test was used to obtain the significance threshold for the AUC (100 times of permutations for each subject), where trial labels were randomly permuted and LOO was carried out. The null distribution of AUC values was generated and a threshold of $p < 0.01$ was used. Details of the STV analysis are illustrated in the Fig. S5.

#### Structural and Functional MRI Analysis

##### Data preprocessing

The fMRI data were processed using FSL (V6.0) [16]. Briefly, motion correction was performed using rigid-body registrations on all the volumes in reference to the middle time point volume [9]. Then, slice timing correction was carried out with Fourier-space time-series phase-shifting. The non-brain tissues were removed using BET [15]. After that, grand-mean intensity normalization was applied by scaling the entire 4D data with a multiplicative factor. Lastly, a high-pass filtering (Gaussian-weighted least-squares straight line fitting, cut-off frequency 0.01 Hz) was carried out. In the spatial normalization, the middle time point EPI volume head image of the fMRI data was rigid registered (cost function as correlation) to the high resolution T2*w head image. Then, the T2*w head image was rigid registered (Boundary-based method) [7] to the T1w head image. Lastly, the subject's T1w brain image was initially affine transformed and then non-linearly registered to the MNI152 (the nonlinear 6th generation atlas from

FSL) brain image using FLIRT [9] and FNIRT [3] from the FSL software package. The structural T1w images were processed using FreeSurfer pipeline [5], which resulted in brain tissue segmentations and surfaces reconstruction. The FreeSurfer segmentations of the T1w images include gray matter, white matter, lateral ventricles, brain masks, etc.

#### STV EEG-informed fMRI analysis

We fit a GLM for each EEG STV time window. First-level GLM was performed independently on each voxel using multiple regression with five variables of interest. The regressors included: 1) event-related regressors with unmodulated height, and both onset and duration matched to the presence of the stimulus (one each for targets and standards); 2) RT variability regressor with unmodulated height, onset matched to the stimulus onset, and duration matched to the RT of the trial (orthogonalized with respect to the targets event-related regressor); 3) EEG STV regressors with height parametrically modulated using the demeaned EEG STV discriminating component, onset set to the time of interest $\tau$, and duration fixed to 100 ms (one each for targets and standards, orthogonalized with respect to the corresponding event-related regressor, oddball EEG STV regressor was also orthogonalized to RT regressor); 4) confounds (motion parameters, temporal derivatives of the variables of interest). At each time window $\tau$, the demeaned output of the logistic regression classifier, as the EEG STV discriminating component $\tilde{d_i}(\tau) = d_i(\tau) - d_{mean}(\tau)$ was used to modulate the height of the EEG STV regressor boxcar function. All regressors were convolved with canonical Double-Gamma hemodynamic response function (HRF). The preprocessed fMRI data were spatially smoothed with a Gaussian kernel of FWHM 5 mm, then were fit with the GLM resulting to five different statistical parametric maps which will be warped onto a standard space (i.e. MNI152) to be able to perform group-level statistical analysis. For group-level statistical inference, we used FMRIB's Local Analysis of Mixed Effects (FLAME) from the FSL software package, where a mixed effect model was carried out, and the group-level statistical parametric maps were thresholded ($z > 2.3$, corrected cluster significance threshold of $p = 0.05$; Gaussian random field).

#### Salience processing node definition

The oddball EEG STV related GLM statistical maps at each time of interest from the STV EEG-informed fMRI analysis were extracted for defining the nodes associated with the processing of oddball trials. At the peak voxel of each group-level significant cluster, a spherical ROI (10 mm radius) centered on the voxel was generated. Centroid of peak locations was used for regions involved in more than one temporal windows (i.e. lS1, rSPL, and mPFC-SMA). After excluding one cluster with the peak voxel outside of the brain mask, ten ROIs were included for the subsequent analyses. The neuroanatomical localizations of the nodes were referred to the HCP-MMP brain parcellation [6]. Specifically, to alleviate inter-subject variability in the cortical surface reconstruction, instead of mapping the HCP-MMP cortex parcellation to the volumetric space, the ROI masks in each subject's native structural space were projected to the subject's cortical surface. After surface-based spatial normalization to the FreeSurfer fsaverage template cortical surface [4], a majority vote was caried out across subjects to obtain the group-level ROI surface areas, which will be compared to the HCP-MMP parcellation.

#### SN, DMN, and DAN node definition

The locations of the nodes in the salience network (SN), default mode network (DMN), and dorsal attention network (DAN) were defined with the HCP-MMP atlas. The SN comprised three nodes: right and left AI (each side includes area anterior agranular insular complex and middle insular area) and dorsal ACC (includes area dorsal 32, anterior 32 prime, and p32 prime; details of area naming in [6]). The DMN comprised five nodes: posterior cingulate cortex (includes RSC, 23d, 23c, d23ab, v23ab, 31a, 31pv, 31pd, POS1, POS2, 7m, DVT, ProS), precuneus (includes PCV), right and left angular gyrus (each side includes PGi, PGs, and PGp), and medial prefrontal cortex (includes 8BM, 9m, 10r, and 10v). The DAN comprised four nodes: right and left SPL (each side includes LIPv, LIPd, VIP, AIP, MIP, 7PC, 7AL, 7Am, 7PL, 7Pm, IP0, IP1, and IP2), and right and left frontal eye fields. The HCP-MMP atlas was transformed to each subject's cortical surface through surface-based registration using FreeSurfer. Then, the selected ROI surface areas were projected into the volumetric space. The ROI masks were warped into the MNI152 template space with the previously estimated registration parameters. A majority vote was caried out across subjects to obtain the group-level ROI masks. After thresholding the ROI masks (0.5 overlap rate), we extracted a weighted center of gravity for each ROI region. Similar to the previous analysis, a spherical ROI (10 mm radius) centered on that voxel was generated. Finally, these twelve nodes were used for the subsequent state-space modeling. Details of the SN, DMN, and DAN nodes are illustrated in the Fig. S6.

##### Functional connectivity analysis

To control for physiological and motion-related noise, we regressed out motion-related nuisance signals from the preprocessed fMRI data (motion parameters included six standard head motion parameters, their temporal derivatives, and the squares of the above twelve motion parameters), as well as the signals in the left and right hemisphere white matter and lateral ventricles. To circumvent the error in spatial normalization and smoothing, we extracted the BOLD time series of the ROIs in each subject's native functional EPI space. The ROI masks were transformed and warped into the subjects' EPI functional space with the estimated registration parameters. For each ROI, only the gray matter BOLD signal was extracted by computing the intersection between the ROI masks and the gray matter mask. And we extracted a single BOLD time series from each ROI by averaging the time series of all the voxels within the intersected mask. No spatial smoothing was carried out. Before computing the FC, to remove the effects of the task, we regressed out task-evoked activations (both standard and target) from the ROIs BOLD signals. In the seed-based FC analysis for network localization, we used a mixed effects model. The first-level seed-based FC was calculated based on the Pearson correlation between the time series of each ROI and the time series of all the voxels in the brain. Then, each subject's FC map was transformed into z-score with Fisher's Z transformation. And the FC z-score map was thresholded at $p < 0.01$. In the group-level, one sample student's t-test was performed to obtain the significant seed-based FC map of each ROI ($p < 0.001$ uncorrected). Similarly, in the node-by-node FC analysis, the first-level FC was computed based on the Pearson correlation across the time series of ROIs. Then, the FC matrix was transformed into z-score, and carried out to the group-level one sample student's t-test to obtain the significant FC matrix across ROIs ($p < 0.05$ uncorrected).

##### Locus coeruleus functional connectivity analysis

To assess functional connectivity between the locus coeruleus (LC) and salience processing nodes, we localized the LC in each subject's functional space with a predefined LC atlas [11] and the subject's TSE image, and then the LC BOLD signal was extracted. Specifically, we first performed a rough localization by estimating the spatial range of the LC location in each subject's structural T1-weighted space, where a statistical criterion was used along with the TSE image intensity spatial distribution and the LC atlas. Then, the TSE image intensity within the estimated range was transformed into the subject's functional EPI space for a precise localization of the LC. The LC BOLD signal was extracted by averaging the voxel-wise BOLD time series, weighted by the TSE image intensities. Details are in [8]. Functional connectivity was computed with Pearson correlation between the BOLD signals extracted from LC and salience processing nodes (mixed effect, $p < 0.05$ uncorrected).

#### Effective Connectivity Analysis

##### State-space modeling of the latent neural activity

A state-space model was used to infer the latent brain dynamics across the nodes. To infer the activity in the EEG source space, a volumetric source model was used, where EEG sources (denoted as $x_t$ at time $t$) are assumed to be uniformly distributed on a 3D grid inside the brain. Given the observations of the scalp EEG measurements $y_t$, a linear EEG forward model was used: $y_t = Lx_t + e_t$, where $L$ is the lead field matrix and $e_t$ is a Gaussian channel noise vector. We defined the latent state variables as $s_t$, where $x_t = Gs_t + \epsilon_t$, $G$ is a binary indicator matrix, and $\epsilon_t$ is a Gaussian noise vector. The influences across the latent states of neural dynamics were modeled as a multivariate autoregressive (MVAR) process as $s_t = As_{t-1} + \sum_{k=1}^{K} B^k m_t^k s_{t-1} + Du_t + \omega_t$, where $A$ is the intrinsic connectivity matrix, $B^k$ is the $k^{th}(k = 1, 2, ...K)$ modulatory connectivity matrix, $m_t^k$ is the modulatory input, $u_t$ is the external input, $D$ is a diagonal matrix denotes the strength of $u_t$, and $\omega_t$ is a Gaussian state noise vector. As for the model inference, the mean-field variational Bayesian approximation was used to make inference on the posterior distributions of the latent space variables and model parameters, during which the evidence lower bound (ELBO) was maximized. The model is described in detail elsewhere [17]. In this study, we first used the ten salience processing nodes defined from the STV EEG-informed fMRI analysis results. Then, another model was fit with the twelve nodes of SN, DMN, and DAN defined from the HCP-MMP atlas. The ROIs were transformed into each subject's native structural space. And each subject's T1w image was used to construct the volumetric head model and the source model using the FieldTrip toolbox [13]. The state-space model was fit to each run separately. The modulatory inputs were modeled as a unit height boxcar function with a sequence of events (one each for targets and standards). Here, the ELBO was also used to quantify the quality of the model fitting. For the data of the runs failed in model fitting, we excluded those runs from subsequent analyses. Specifically, 11 out of 90 runs failed in the model fitting with salience processing nodes, and 7 out of 90 runs failed in the model fitting with SN, DMN, and DAN nodes.

##### Bayesian parameter averaging

The estimated posterior distributions of model parameters in the first-level analysis were summarized for group-level Bayesian posterior inference using Bayesian parameter averaging (BPA) [12], which computes a group-level joint posterior probability for the effect of interest. Compared to the conventional statistical null hypothesis significance tests (NHST), the Bayesian posterior inference is considered more robust as the influence from each run is weighted by the precision. The significant group-mean EC was estimated, where a posterior probability criterion of $\alpha < 0.05$ (Bonferroni correction) was used to threshold the group-level posterior distribution for the inference of significant EC.

#### Pupillometry Analysis

##### Data preprocessing and epoching

The preprocessing pipeline was adapted from the approach in [18]. Firstly, blink detection was performed with Eyelink software, and then the blinks were padded by 150 ms and linearly interpolated. Additional blinks were further removed with a peak detection algorithm. We computed pupil diameter from the pupil area data, and then the pupil diameter time series were filtered with a bandpass second-order Butterworth filter (0.01 Hz to 10 Hz). Then, the pupil diameter data of each run were z-scored independently, and down-sampled to 500 Hz (the same sampling rate of the preprocessed EEG data). The preprocessed pupil diameter data were epoched from 500 ms before stimulus to 2000 ms following the stimulus. Two pupil diamster measurements were examed including the prestimulus baseline pupil diameter (BPD) and the TEPR. The BPD was defined as the pupil diameter at stimulus onset time, and TEPR was defined as the maximum percentage deviation from BPD within each epoch.

##### Brain and pupil correlation analysis

To assess the relationship between the network-level interactions and pupillary response, we characterized the network-level connectivity in the positive and negative connections. Specifically, we defined positive and negative network interaction strength as the sum of all positive and negative connection parameters from one network node set to the other network node set (or to itself as self-connection network strength), respectively. And we also computed the mean TEPR across all the oddball trials for each subject. Therefore, we performed a correlation analysis across all subjects between the network interaction strength and the mean TEPR (Pearson correlation). As a control analysis, we also regressed out the median RT and the mean ELBO from the mean TEPR and network interaction strength, respectively, before performing the same analysis across subjects.

### Results

#### Relationship between Effective Connectivity of Networks and Pupillary Response

We evaluated the association between the late-to-early positive network strength and TEPR by computing the Pearson correlation between them. The significance level was set as $\alpha < 0.05$ with Bonferroni correction. For the positive connections, whilst the late-to-early network strength showed a significant correlation with TEPR (r = 0.6352, p = 0.0035), the other network connectivity strength did not show any statistical significant relationship with TEPR (Early-to-early: r = 0.4290, p = 0.0668; Early-to-middle: r = 0.3708, p = 0.1181; Early-to-late: r = 0.0916, p = 0.7091; Middle-to-early: r = 0.1163, p = 0.6354; Middle-to-middle: r = 0.1219, p = 0.6191; Middle-to-late: r = 0.1058, p = 0.6665; Late-to-middle: r = 0.1404, p = 0.5665; Late-to-late: r = -0.1568, p = 0.5214). For the negative connections, none of these network connections strength showed significant correlation with TEPR.

##### Functional Connectivity between Locus Coeruleus and Salience Processing Nodes

The LC showed significant functional connectivity with lSPL, lS1 and mPFC-SMA, however, there were no significant results between LC and the other salience processing nodes (rM1: t = 0.73, p = 0.476; rV2: t = -1.15, p = 0.267; rSPL: t = 1.48, p = 0.156; lIPL: t = -0.45, p = 0.660; lOFC: t = -0.56, p = 0.579; rOFC-rIFC: t = -1.92, p = 0.071; lAuditory and Frontal operculum: t = 0.96, p = 0.351).

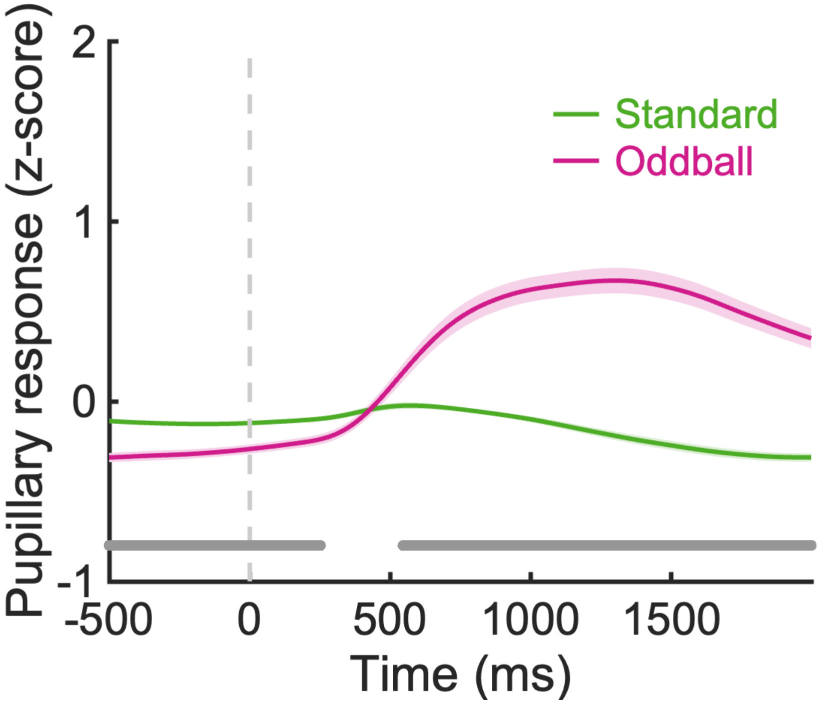


**Figure S1.** Stimuli-locked pupillary response. The z-scored pupil diameter fluctuations from 500 ms before the stimulus to 2000 ms following the stimulus were averaged across subjects for the oddball (pink) and standard (green) stimuli. The shaded bands represent standard error, and the bottom gray line indicates significant difference (Student's t-test, $p < 0.001$) between the pupil diameter evoked by the oddball and standard stimuli.

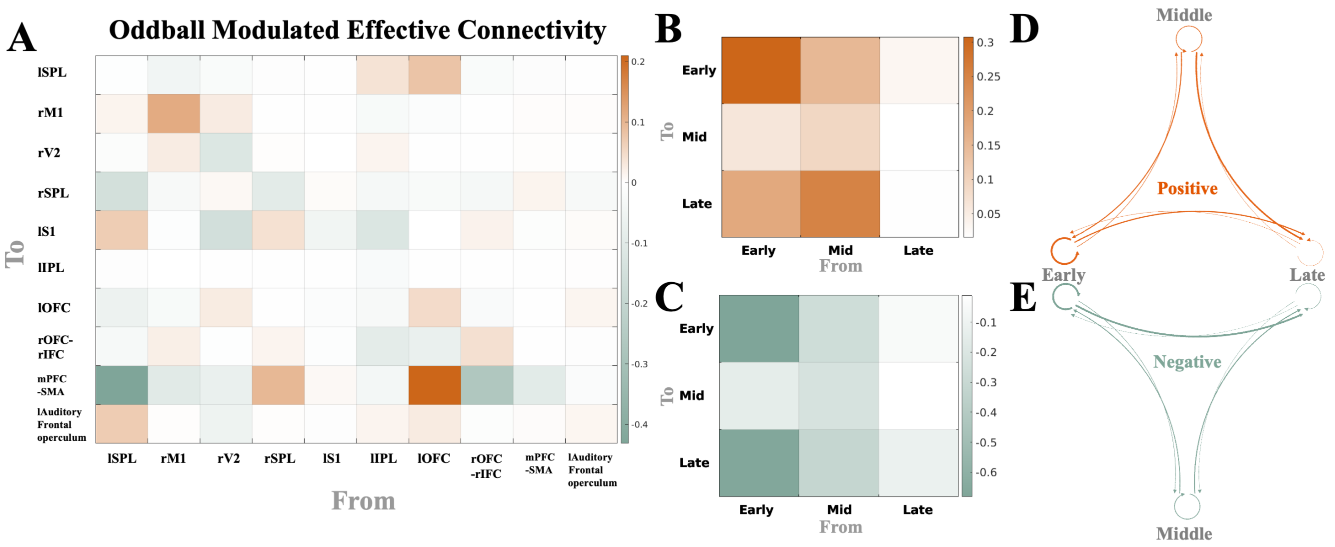


**Figure S2.** Group-level mean effective connectivity modulated by the oddball stimuli, with node-wise effective connectivity matrix (A), network-wise positive (B and D) and negative (C and E) effective connectivity. The group-level network-wise mean effective connectivity was computed as the sum of all positive (or negative) group-level connection parameters from one network node set to the other network node set. Please be noted the results here reflect mean group effect. The thickness of the connecting arrows in D and E corresponds to the strength of effective connectivity. The orange and blue color represents positive and negative effective connectivity, respectively.

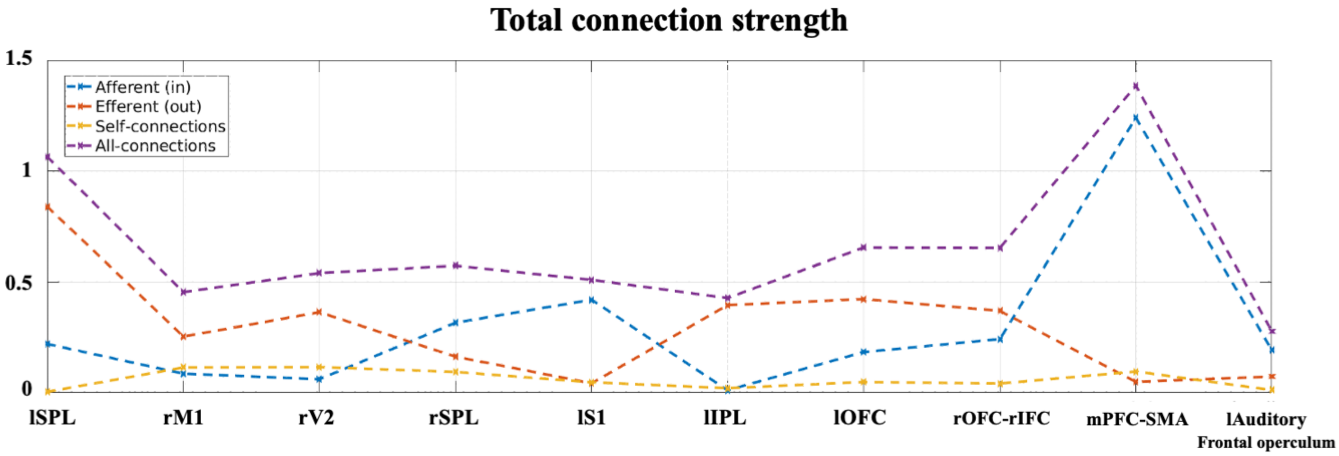


**Figure S3.** Total connection strength of each salience processing node. All the unsigned connection parameters (efferent, afferent and self-connection) associated with the node were summed up to compute the total connection strength. The results suggest that the lSPL and mPFC-SMA have the strongest total connection strength, indicating their roles as hubs in the processing of salience stimuli.

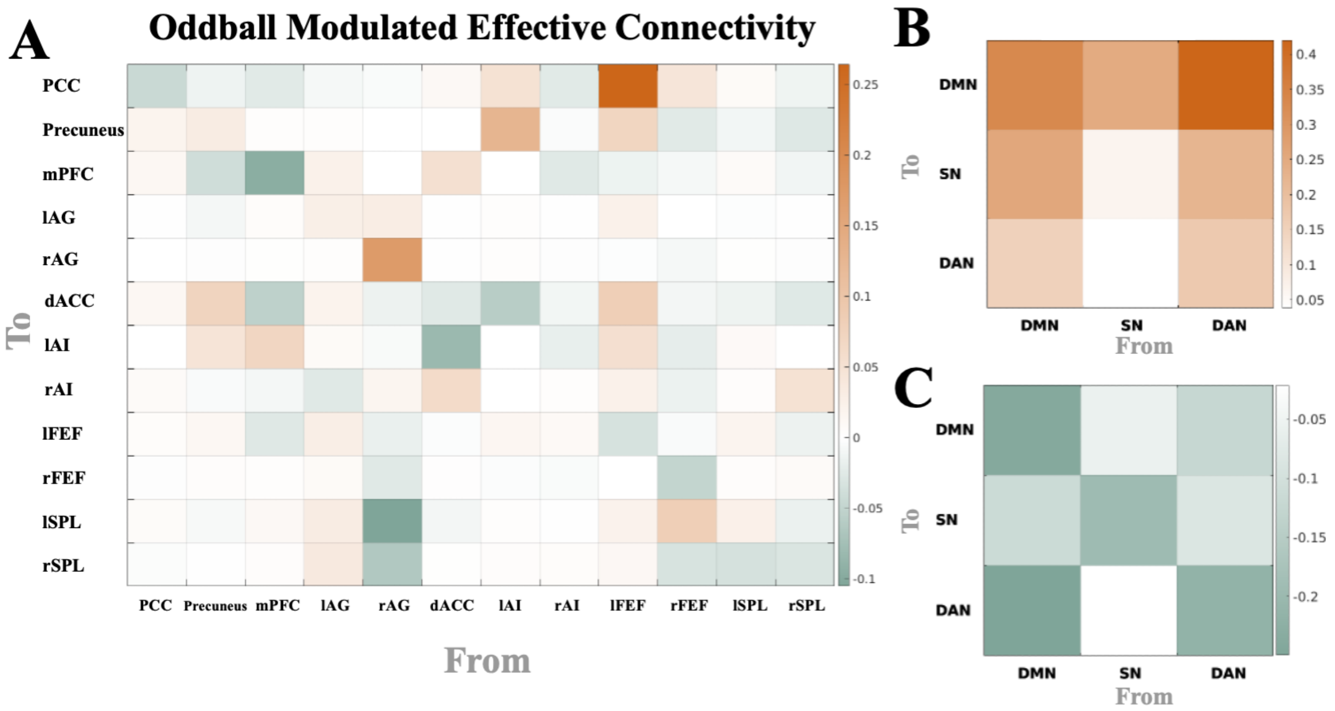


**Figure S4.** Group-level mean effective connectivity modulated by the oddball stimuli across DMN, SN and DAN, with node-wise effective connectivity matrix (A), network-wise positive (B) and negative (C) effective connectivity. The orange and blue color represents positive and negative effective connectivity, respectively.

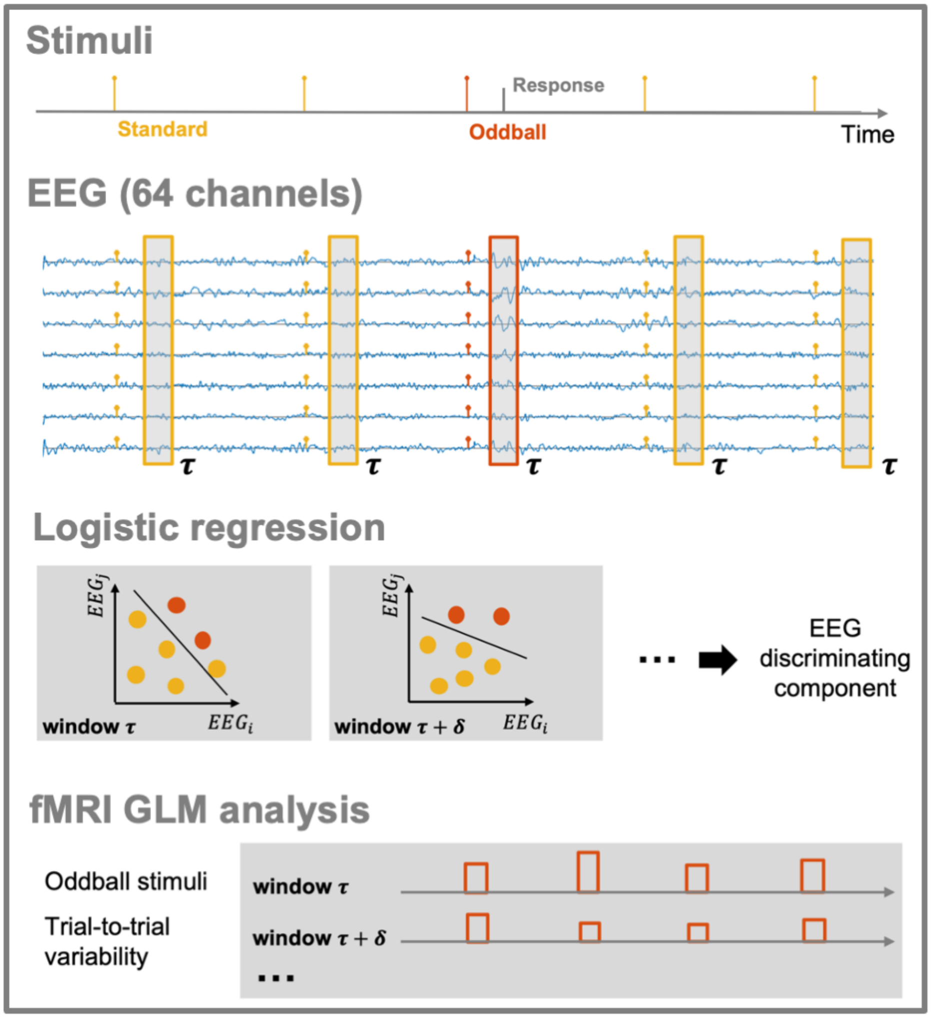


**Figure S5.** Auditory oddball paradigm, single-trial analysis, and single-trial variability EEG-informed fMRI analysis. For each temporal window $\tau$, we applied a single-trial analysis with the extracted EEG data in the windows from all the trials, where a logistic regressor was trained to learn a weight matrix w maximally discriminating the target vs standard trials. From the weighting on the EEG channels with matrix w, a EEG discriminating component was computed as a low-dimensional representation of the EEG data. For example, two EEG sensors (channel i and j) were illustrated in the figure with a hyperplane discriminating target (red dots) and standard (yellow dots) trials. Similarly, single-trial analysis was applied to all other temporal windows spanning the trial independently with a sliding window approach (step size as $\delta$). The EEG discriminating component at each temporal window was used to modulate regressors in a general linear model (GLM) to predict fMRI BOLD response (convolved with the canonical haemodynamic response function along with other regressors). The GLM analysis was applied with each temporal window independently.

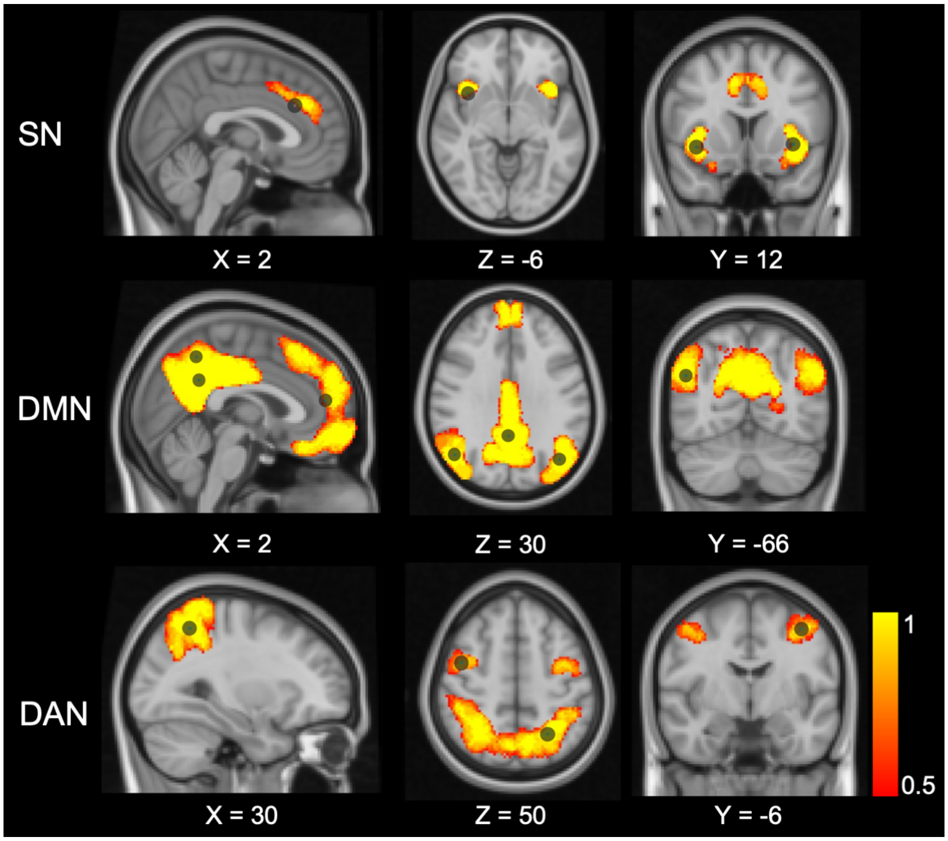


**Figure S6.** Network nodes definition with HCP-MMP atlas. The nodes (circles) of the SN, DMN, and DAN are overlaid with the selected network areas from the HCP-MMP atlas and the MNI152 brain image. The group-level region of interest masks (illustrated as spatial distribution maps) were obtained from majority vote across subjects.

### References


1. R. Abreu, A. Leal, and P. Figueiredo. EEG-informed fMRI: a review of data analysis methods. *Frontiers in Human Neuroscience*, 12:29, 2018.
2. P. J. Allen, O. Josephs, and R. Turner. A method for removing imaging artifact from continuous EEG recorded during functional MRI. *NeuroImage*, 12(2):230–239, 2000.
3. J. L. Andersson, M. Jenkinson, S. Smith, et al. Non-linear registration, aka spatial normalisation FMRIB technical report TR07JA2. *FMRIB Analysis Group of the University of Oxford*, 2(1):e21, 2007.
4. B. Fischl, M. I. Sereno, R. B. Tootell, and A. M. Dale. High-resolution intersubject averaging and a coordinate system for the cortical surface. *Human Brain Mapping*, 8(4):272–284, 1999.
5. B. Fischl, A. Van Der Kouwe, C. Destrieux, E. Halgren, F. Ségonne, D. H. Salat, E. Busa, L. J. Seidman, J. Goldstein, D. Kennedy, et al. Automatically parcellating the human cerebral cortex. *Cerebral Cortex*, 14(1):11–22, 2004.
6. M. F. Glasser, T. S. Coalson, E. C. Robinson, C. D. Hacker, J. Harwell, E. Yacoub, K. Ugurbil, J. Andersson, C. F. Beckmann, M. Jenkinson, et al. A multi-modal parcellation of human cerebral cortex. *Nature*, 536(7615):171–178, 2016.
7. D. N. Greve and B. Fischl. Accurate and robust brain image alignment using boundary-based registration. *NeuroImage*, 48(1):63–72, 2009.
8. H. He, L. Hong, and P. Sajda. An automatic and subject-specific method for locus coeruleus localization and BOLD activity extraction. In *Proc. Intl. Soc. Mag. Reson. Med.*, 2021.
9. M. Jenkinson, P. Bannister, M. Brady, and S. Smith. Improved optimization for the robust and accurate linear registration and motion correction of brain images. *NeuroImage*, 17(2):825–841, 2002.
10. T.-P. Jung, S. Makeig, M. Westerfield, J. Townsend, E. Courchesne, and T. J. Sejnowski. Removal of eye activity artifacts from visual event-related potentials in normal and clinical subjects. *Clinical Neurophysiology*, 111(10):1745–1758, 2000.
11. N. I. Keren, C. T. Lozar, K. C. Harris, P. S. Morgan, and M. A. Eckert. In vivo mapping of the human locus coeruleus. *NeuroImage*, 47(4):1261–1267, 2009.
12. J. Neumann and G. Lohmann. Bayesian second-level analysis of functional magnetic resonance images. *NeuroImage*, 20(2):1346–1355, 2003.
13. R. Oostenveld, P. Fries, E. Maris, and J.-M. Schoffelen. FieldTrip: open source software for advanced analysis of MEG, EEG, and invasive electrophysiological data. *Computational Intelligence and Neuroscience*, 2011, 2011.
14. L. C. Parra, C. D. Spence, A. D. Gerson, and P. Sajda. Recipes for the linear analysis of EEG. *NeuroImage*, 28(2):326–341, 2005.
15. S. M. Smith. Fast robust automated brain extraction. *Human Brain Mapping*, 17(3):143–155, 2002.
16. S. M. Smith, M. Jenkinson, M. W. Woolrich, C. F. Beckmann, T. E. Behrens, H. Johansen-Berg, P. R. Bannister, M. De Luca, I. Drobnjak, D. E. Flitney, et al. Advances in functional and structural MR image analysis and implementation as FSL. *NeuroImage*, 23:S208–S219, 2004.
17. T. Tu, J. Paisley, S. Haufe, and P. Sajda. A state-space model for inferring effective connectivity of latent neural dynamics from simultaneous EEG/fMRI. *Advances in Neural Information Processing Systems*, 32:4662–4671, 2019.
18. A. E. Urai, A. Braun, and T. H. Donner. Pupil-linked arousal is driven by decision uncertainty and alters serial choice bias. *Nature Communications*, 8(1):1–11, 2017.
